# A physiologically-based digital twin for alcohol consumption – predicting real-life drinking responses and long-term plasma PEth

**DOI:** 10.1101/2023.08.18.553836

**Authors:** Henrik Podéus, Christian Simonsson, Patrik Nasr, Mattias Ekstedt, Stergios Kechagias, Peter Lundberg, William Lövfors, Gunnar Cedersund

## Abstract

Alcohol consumption is associated with a wide variety of preventable health complications and is a major risk factor for all-cause mortality in the age group 15-47 years. To reduce dangerous drinking behavior, eHealth applications have shown promise. A particularly interesting potential lies in the combination of eHealth apps with mathematical models. However, existing mathematical models do not consider real-life situations, such as combined intake of meals and beverages, and do not connect drinking to clinical markers, such as *phosphatidylethanol* (PEth). Herein, we present such a model which can simulate real-life situations and connect drinking to long-term markers. The new model can accurately describe both estimation data according to a χ^2^-test (187.0 < T_χ2_ = 226.4) and independent validation data (70.8 < T_χ2_ =93.5). The model can also be personalized using anthropometric data from a specific individual and can thus be used as a physiologically-based digital twin. This twin is also able to connect short-term consumption of alcohol to the long-term dynamics of PEth levels in the blood, a clinical biomarker of alcohol consumption. Here we illustrate how connecting short-term consumption to long-term markers allows for a new way to determine patient alcohol consumption from measured PEth levels. An additional use case of the twin could include the combined evaluation of patient-reported AUDIT forms and measured PEth levels. Finally, we integrated the new model into an eHealth application, which could help guide individual users or clinicians to help reduce dangerous drinking.

## Introduction

Alcohol consumption causes around five percent of all deaths worldwide, in addition to contributing to numerous societal health issues (1). Long-term alcohol consumption is associated with an increased risk of chronic liver disease, *hepatocellular carcinoma* (HCC), and other malignancies (2,3). Additionally, how one consumes alcohol, *i.e.,* one’s personal drinking habits (4), might contribute to both short-term and long-term effects on personal health. One common type of drinking habit is ‘binge drinking’ (heavy episodic drinking), which is more prevalent in adolescents and young adults (5,6). Binge drinking can cause major health problems, contributing to both acute injuries (*e.g.,* accidents) and long-term negative effects (7). Furthermore, during ‘heavy drinking’, the ability to judge your own intoxication level is decreased (8). For these and other reasons, it is important to determine and control the amount of alcohol consumed and highlight the long-term effects. When it comes to reducing harmful alcohol use, as well as improving care, eHealth applications can be useful.

Various types of eHealth applications and digital tools have shown promise in affecting alcohol consumption patterns in individuals with unhealthy alcohol use or *alcohol use disorder* (AUD) (9–16). For example, in a behavioral intervention study by Bendtsen et al. (17) a digital intervention produced self-reported changes in alcohol consumption among online help-seekers in the general population. Additionally, some eHealth applications and interventions include *estimated blood alcohol concentration* (eBAC) calculations (18–21). For example, the eBAC-based application “*Promillekollen”* was used in a study by Berman et al. (18). In this study, a group of university students, with hazardous alcohol use, was shown to reduce their number of ‘binge drinking’ sessions by using the application. Apart from these examples, eHealth applications could potentially be developed for a variety of other use cases: e.g. clinical diagnosis, prognosis, and patient education. Many of those use cases would require a long-term predictive ability, which takes into account real-life drinking patterns, and predicts long-term alcohol markers, such as *phosphatidylethanol* (PEth) (Fig. 1a) (22).

**Figure 1:**
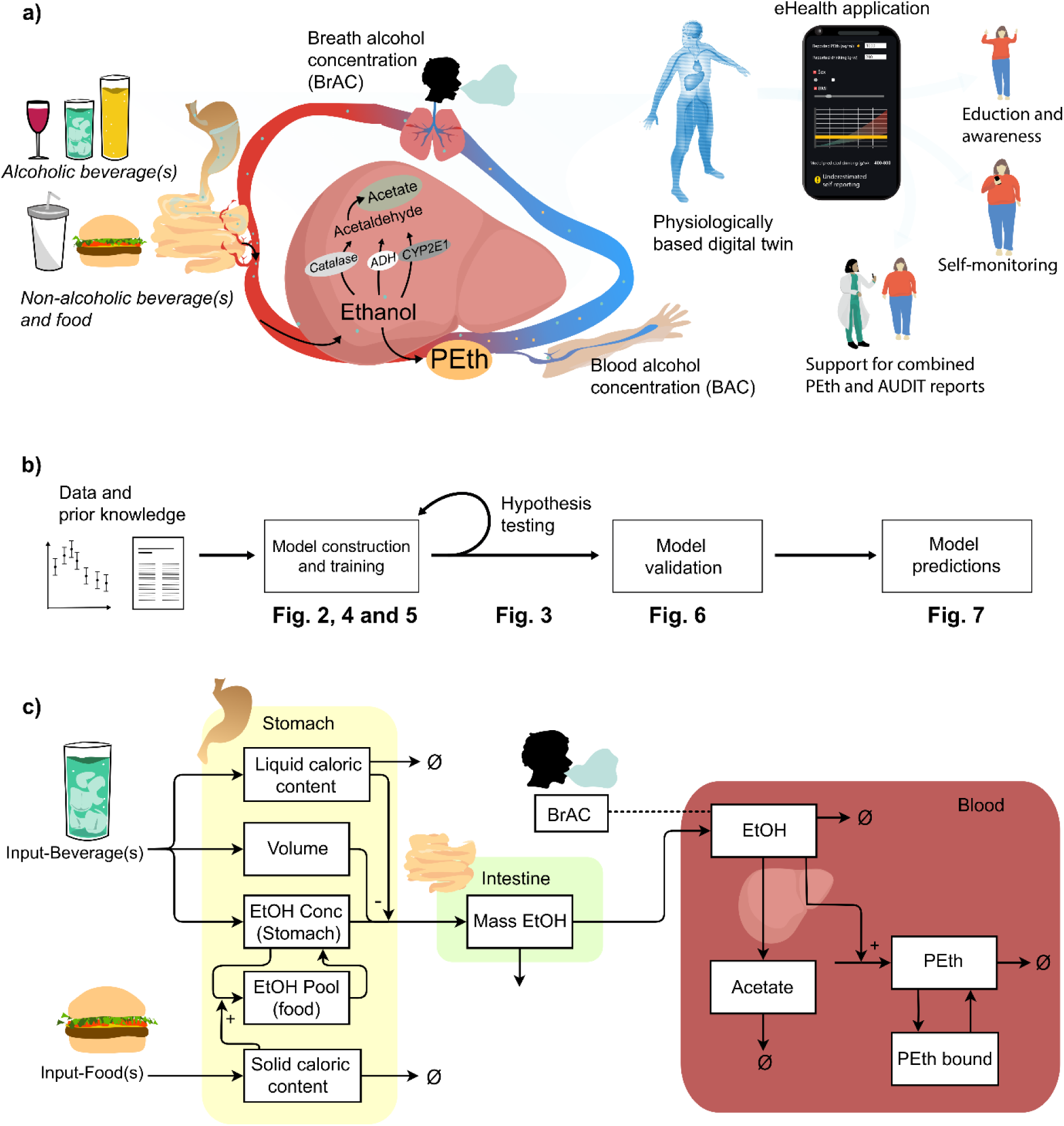
Overview of the physiological processes and use cases, modelling framework and mathematical model representation. **a)** Short overview of the physiological process that the model describes. Ethanol enters the body via the stomach, where already a small amount can enter plasma. Via the stomach emptying the ethanol enters the intestine. Here, most of the ethanol is taken up via absorption. Most of the ethanol is metabolized in the liver, and a small amount is excreted via renal pathways. In the liver, ethanol is converted into acetaldehyde via three oxidative pathways governed by the enzymes: *Alcohol dehydrogenases* (ADH), catalase, and *cytochrome P450 2E1* (CYP2E1). Acetaldehyde is further converted into acetate and then acetyl-COA. There also exist non-oxidative pathways, responsible for a miniscule amount of ethanol breakdown, e.g., into *phosphatidylethanol* (PEth). Following, the *blood alcohol concentration* (BAC), or the *breath alcohol concentration* (BrAC) is measured. These physiological processes can be described using a mathematical model, a physiologically-based digital twin. The digital twin can be used for several use cases such as: for education and awareness, in self-reporting and monitoring of alcohol consumption, and as a tool to support the combination of AUDIT and PEth reports. **b)** Schematic over the modelling approach. **c)** Schematic showing the model structure, for a detailed description see material and methods.

Measuring PEth has been shown to be useful for characterizing actual alcohol consumption (23,24). Currently, alcohol consumption is measured using self-report forms, e.g., *alcohol use disorders identification test* (AUDIT) (25). However, there are serious indications of inconsistencies between self-reported alcohol consumption, and long-term biomarkers in a significant portion of AUD patients (26). A step towards increasing the accuracy of self-reporting forms and developing future screening methods could be to link every instance of alcohol consumption to, e.g., a long-term PEth trajectory. Thus, an eHealth solution, connecting long-term alcohol markers to short-term drinking behavior, would present an option for estimating previous drinking behavior from measured PEth levels. However, for accurate estimates of previous drinking behavior based on PEth levels, one needs a model that connects blood alcohol concentration (BAC) and PEth.

There are models that combine BAC and PEth, but their usefulness is limited, since they do not describe real-life drinking scenarios. A simple and widely used model for BAC estimations is the Widmark equation (27). The simplicity of the Widmark equation implies that it cannot describe many real-life drinking scenarios. Such scenarios include e.g. different types of drinks, as well as drinks in combination with food. Because the Widmark equation does not describe gastric emptying, it can neither describe the slowing effect of beverage caloric content on gastric emptying (28–32), nor the ingestion of meals which greatly reduces the BAC following consumption of alcohol (33–37). There are other more advanced models that describe either (i) the gastric emptying (38–40), (ii) detailed BAC profiles of varying detail (41–45), or (iii) the PEth dynamics (46). However, to our knowledge, there does not exist a mathematical model that can describe all these aspects in a single model. Therefore, current models are insufficient for connecting BAC profiles with PEth for real-life drinking patterns.

Herein, we present a physiologically-based model that incorporates several aspects of BAC after alcohol consumption not presented in other models: 1) factors impacting gastric emptying, 2) alcohol interaction with meals, and 3) ethanol break-down into PEth (Fig. 1c). This model establishes the basis for a physiologically-based digital twin of alcohol consumption (47–49).

## Results

The mechanistic model, *i.e.,* the physiologically-based digital twin was trained and validated on various published experimental data (24,28,29,33,37,50–54). An overview of the framework of this study is given in Fig. 1b and an overview of the estimation and validation data is given in the Supplementary Materials “S4 Usage of experimental data”. The model could simultaneously find agreement with all data included in the estimation dataset using one set of parameters (Fig. 2, 4–5). Additionally, the model can predict independent validation data using the same parameter values (Fig. 6). The model agreement to data was evaluated using a χ^2^-test on the residuals (see methods section). The χ^2^-test cutoff value was chosen using a confidence level α = 0.05, with degrees of freedom being the number of data-points included in the estimation dataset (193). For the best model agreement, the χ^2^ test statistic was 187.0 which was lower than the cutoff (T_χ2_ = 226.4). The model prediction of the validation data (using the parameter values that gave the best agreement to estimation data) also passes a χ^2^-test with a confidence level of α = 0.05. The χ^2^ test statistic was 70.8 which was lower than the cutoff (T_χ2_ = 93.5, for 73 degrees of freedom). Furthermore, we used the modelling framework to test different hypotheses for the effect of meals on the plasma ethanol dynamics (Fig. 3). Lastly, we used the model to predict long-term PEth levels as a response to daily drinking for 90 days (Fig. 7).

**Figure 2:**
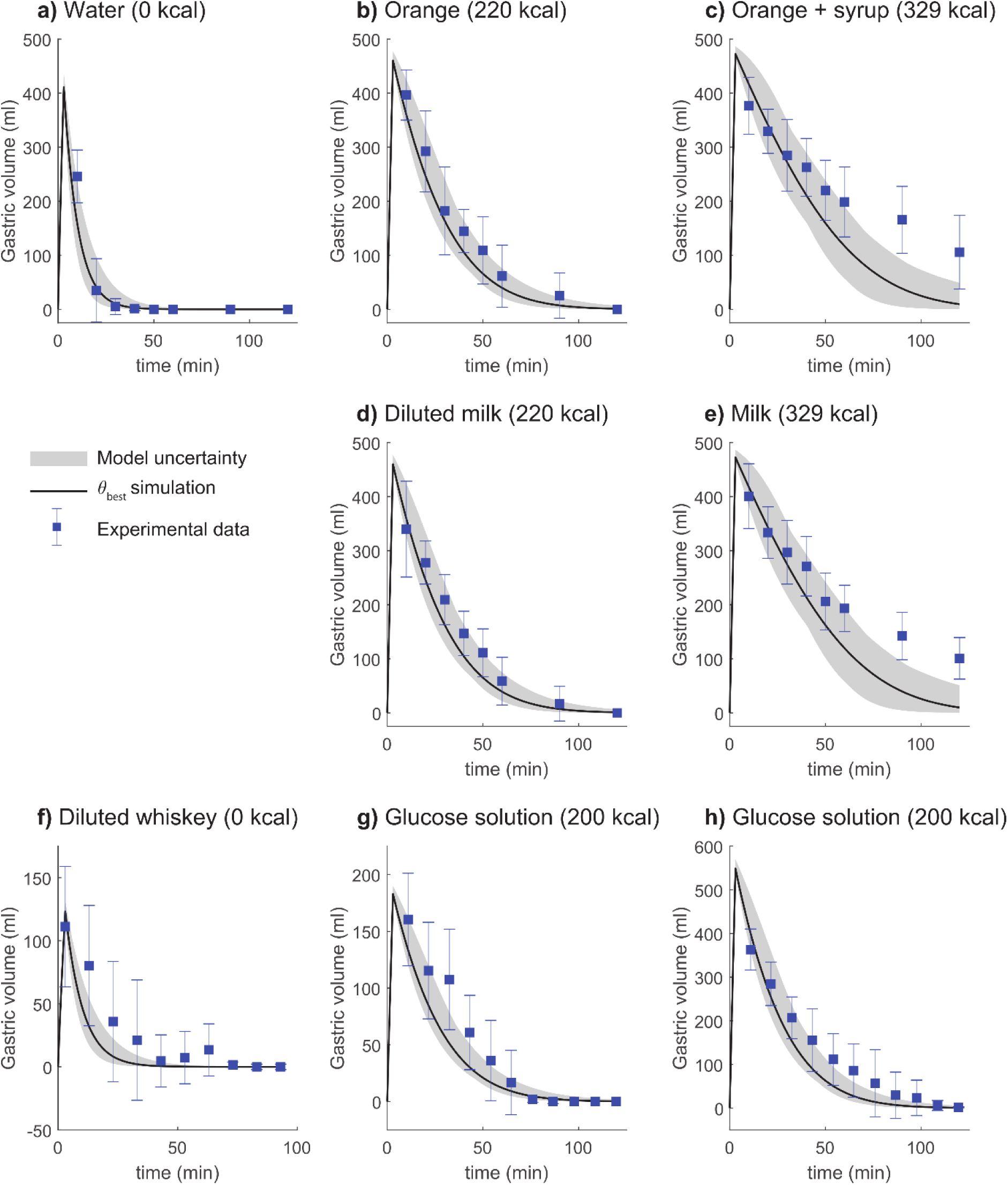
Model agreement to gastric emptying data. The solid line is the best model fit, the shaded area is the model uncertainty, the blue boxes are the mean experimental data value with the error bars indicating the *standard error of the mean* (SEM). The model (solid lines) describes the clearance of different beverages from the stomach (blue markers with error bars). In all experiments, the beverage was consumed over 3 minutes and was constituted of the following: **a)** 500 mL containing 0 kcal, **b)** 500 mL containing 220 kcal, **c)** 500 mL containing 329 kcal, **d)** 500 mL containing 220 kcal, **e)** 500 mL containing 329 kcal, **f)** 150 mL containing 0 non-ethanol kcal, **g)** 200 mL containing 200 kcal, and **h)** 600 mL containing 200 kcal. The important factors in controlling gastric emptying are: 1) the caloric content of the beverage does not show a significant difference (**b** vs. **d** and **c** vs. **e**), and 2) the total caloric content is important rather than the density of calories (**g**, **h**).

### The model can find agreement with gastric emptying data describing different types of interventions

In this section, we will show the model agreement for the gastric emptying module of our model to experimental data from three studies published by Okabe *et al.* (28,29,50). In one of the studies by Okabe *et al.* (28), the study participants drank beverages with the same volume but different caloric content. The data from the study indicates that there is no significant effect of the calorie type on the rate of gastric emptying. This behavior is captured by the model, where the different caloric content shows the same gastric emptying rate (compare Fig. 2b with Fig. 2d, and Fig. 2c with Fig. 2e). Rather, the total caloric content seems to be the driving factor in slowing down the rate of gastric emptying (compare Fig. 2a (0 kcal) with Fig. 2b and d (220 kcal), and Fig. 2c and e (339 kcal)). In the second study by Okabe *et al.* (50), participants ingested an alcoholic whiskey based drink (8.0 v/v%), and the gastric volume was measured, which the model describes (Fig. 2f). In the third study by Okabe *et al*. (29), experimental data indicates that caloric density is not of importance for the gastric emptying, rather the total caloric content is the most significant factor (29). The model can describe this behavior as both volumes are emptied at a similar rate, given the same total caloric content and different caloric densities (Fig. 2g-h). In summary, our gastric emptying module can find agreement with a range of experimental data and captures fundamental mechanisms shown in the data (28,29,50).

### Investigation of the meal effect on plasma ethanol dynamics

The process of the model formulation (Fig. 1b) of the meal effect on plasma ethanol is showcased here (Fig. 3). Four different hypotheses (H1-H4) of the interaction between ethanol and the food were formulated based on known interactions (33–36,55). The hypotheses are illustrated in Fig. 3a, as the following: H1) alcohol is encapsulated by food and released when the food is degraded, H2) the food slows down the passing of liquid to the intestines, H3) in addition to slower passing of liquid to the intestines, gastric alcohol dehydrogenase is present and the activity is upregulated by the food, and H4) processing of the food increases blood flow, and by extension the enzyme effect in the liver. All these hypotheses describe the rate of gastric emptying of foodstuff as reported by Tougas *et al.* (56). These hypotheses were all evaluated against the estimation data (T_χ2_ = 226.4) and only H1 could sufficiently describe data with a χ^2^ test of 187.0 (the χ^2^ test statistic for the other hypothesis was: H2 = 816.41, H3 = 844.49, and H4 = 280.05). Additionally, hypotheses H2-H4 had issues in explaining qualitative behaviors. Here, the model behaviors of H1-H4 are shown with a subset of the estimation data (Fig. 3b), more specifically data presented by Jones et al. on alcoholic drinks in combination with food (33). The model behavior of H2 fails to slow down the release of ethanol from the stomach sufficiently as the simulated BAC overshoots the data. H3 slows down the initial appearance of BAC too much resulting in a peak that is delayed in relation to the peak shown in the data. H4 speeds up the elimination of ethanol in the blood too much, as the clearance of BAC is too fast in the second half of the time series. Only H1 can sufficiently slow down the release of ethanol from the stomach to match the behavior described by the estimation data. Thus, the final model formulation includes the meal effect on plasma ethanol as described by H1.

**Figure 3:**
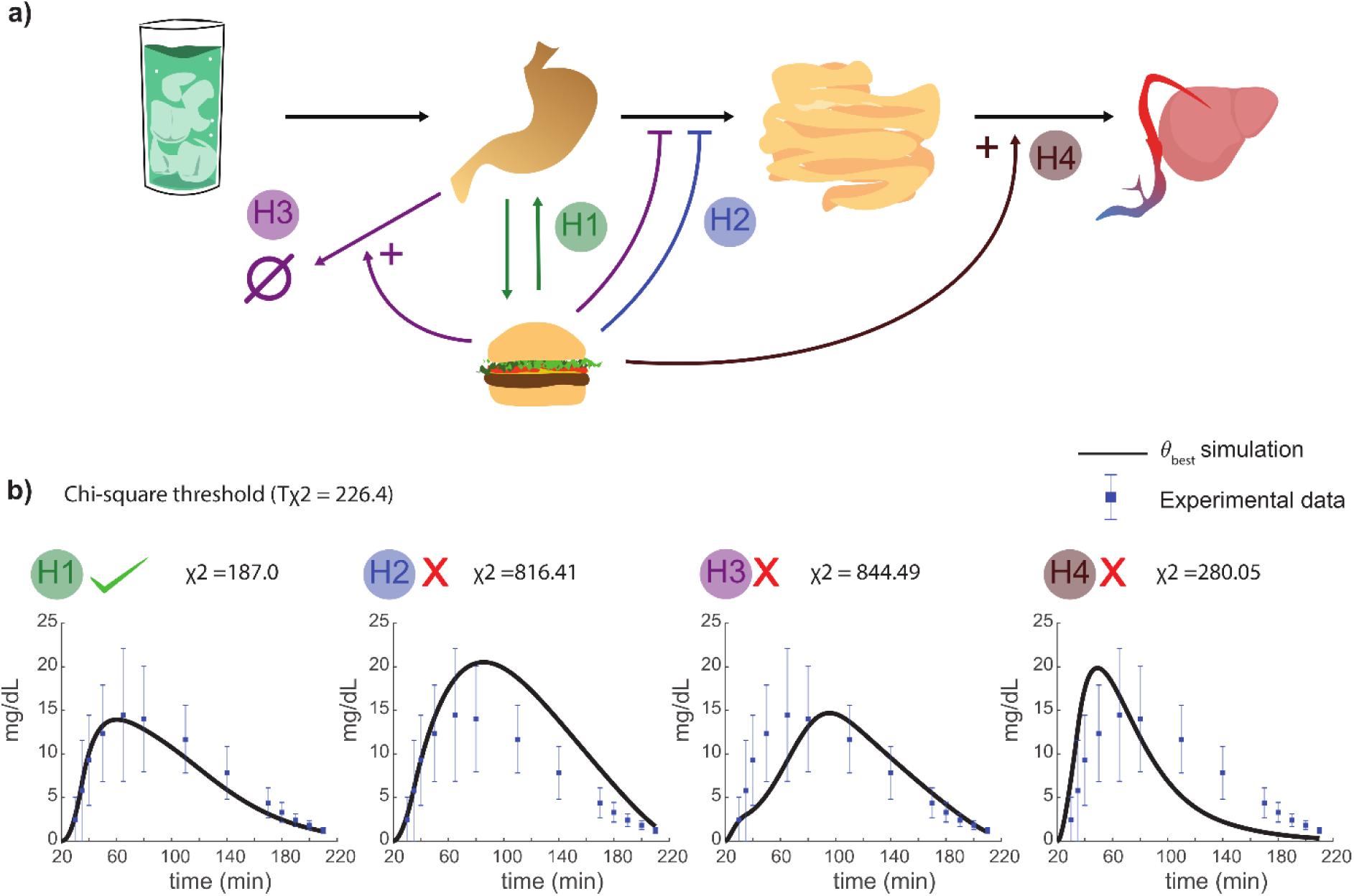
Investigation of the meal effect on plasma ethanol dynamics. **a)** Four different hypotheses were designed that could be explained to describe the meal effect on plasma ethanol dynamics. H1: food can encapsulate ethanol while present in the stomach. While the food is processed, ethanol is released back into the stomach. H2: the contents of the stomach (including the liquid) are held longer due to the food needing to be processed. H3: gastric alcohol dehydrogenase is introduced and is upregulated by food in addition to the contents of the stomach being held longer. H4: the processing of the food recruits more blood to the gastric area and therefore the alcohol transport to the liver is upregulated – increasing the activity of liver alcohol dehydrogenase. **b)** The different model hypothesis was implemented and fitted to the estimation data. Here, model simulation of each respective model (H1-H4) are compared with the experimental data from Jones *et al.* (33), which is one of the included datasets in the estimation data. When evaluating the hypothesis behavior to all the estimation data using a χ2 test statistic – only H1 describes the experimental data sufficiently well.

### The model can describe plasma ethanol data from a wide range of real-life drinking scenarios

In addition to describing the gastric emptying data, the model can simultaneously describe plasma ethanol from a wide variety of experimental studies (33,37,51,52,57). The studies included in the model estimation dataset introduced different types of perturbations. Firstly, the study by Mitchell *et al.* (51) presents the plasma ethanol dynamics after consumption of three different beverages with the same total alcoholic content (Beer: 1 L, 5.1 v/v%; Wine: 0.42 L, 12 v/v%; Spirit: 0.26 L, 20 v/v%). Here, one can observe that increasing alcohol concentration results in a greater BAC amplitude. The model can describe the Mitchell *et al.* data (Fig. 4a-c). Secondly, the model can fit both BAC data with and without food (Fig. g-h, and Fig. a-f respectively). The meal effect results in a decreased amplitude of the BAC profile. This is observed in the data presented by Jones *et al*. and Kechagias *et al*. (33,37), in fasting subjects (Fig. 4d-e) and a fed state (Fig. 4g-h). Lastly, Fig. 4f shows the model agreement to *breath alcohol concentration* (BrAC) data from a study by Javors *et al*. (12). In summary, the model describes several different types of real-life drinking scenarios.

**Figure 4:**
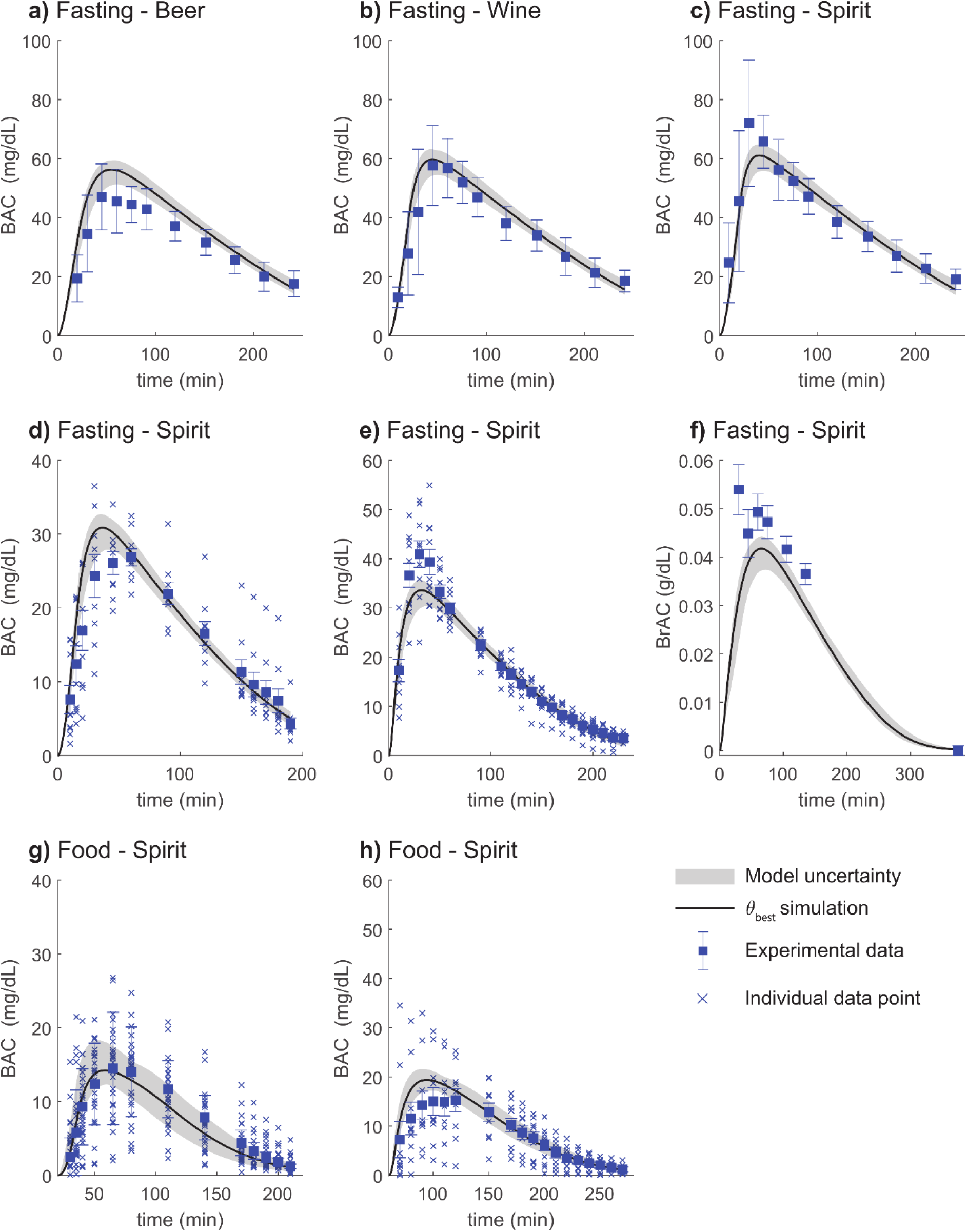
Model agreement to ethanol data for various interventions. The solid line is the best model fit, the shaded area is the model uncertainty, the blue boxes are the mean experimental data value with the error bars indicating the *standard error of the mean* (SEM), and individual data points are marked with a “x”. The alcohol was consumed orally through different beverages in various alterations **a)** 1L of 5.1 v/v % beer (133 kcal) was consumed over 20 min, **b)** 0.42L of 12.5 v/v % wine (56 kcal) was consumed over 20 min, **c)** 0.26L of 20 v/v % spirit blend (43 kcal) was consumed over 20 min, **d)** 0.14L of 20.0 v/v % spirit blend (51 kcal) over 15 min, **e)** 0.15L of 20 v/v % spirit blend (53 kcal) was consumed over 5 min, **f)** 0.71L of 6.5 v/v % spirit blend (247 kcal) was consumed over 15 min, **g)** 0.14L of 20.0 v/v % spirit blend (51 kcal) over 15 min after a meal constituted of 700 kcal, **h)** 0.15L of 20 v/v % spirit blend (53 kcal) was consumed over 5 min after eating a meal constituted of 760 kcal. In **a-f** the beverage was consumed in a fasting state, and in **g**-**h** in combination with food. In **a-e** and **g-h** absolute change of BAC was measured and in **f** absolute change in *breath alcohol concentration* (BrAC) was measured.

### The model can describe products of hepatic ethanol metabolism

Included in the estimation dataset was experimental data describing the plasma concentrations of derivatives of oxidative and non-oxidative ethanol breakdown pathways (Fig. 5a). The data used to represent the oxidative pathways was plasma acetate levels after beverage consumption presented by Sarkola *et al.* The model agreement to the acetate data is shown (Fig. 5b). To represent the non-oxidative breakdown, data describing plasma PEth from a study by Javors *et al.* (52) was included in the estimation dataset. The PEth data included both short-term and long-term data of PEth concentration changes.

**Figure 5:**
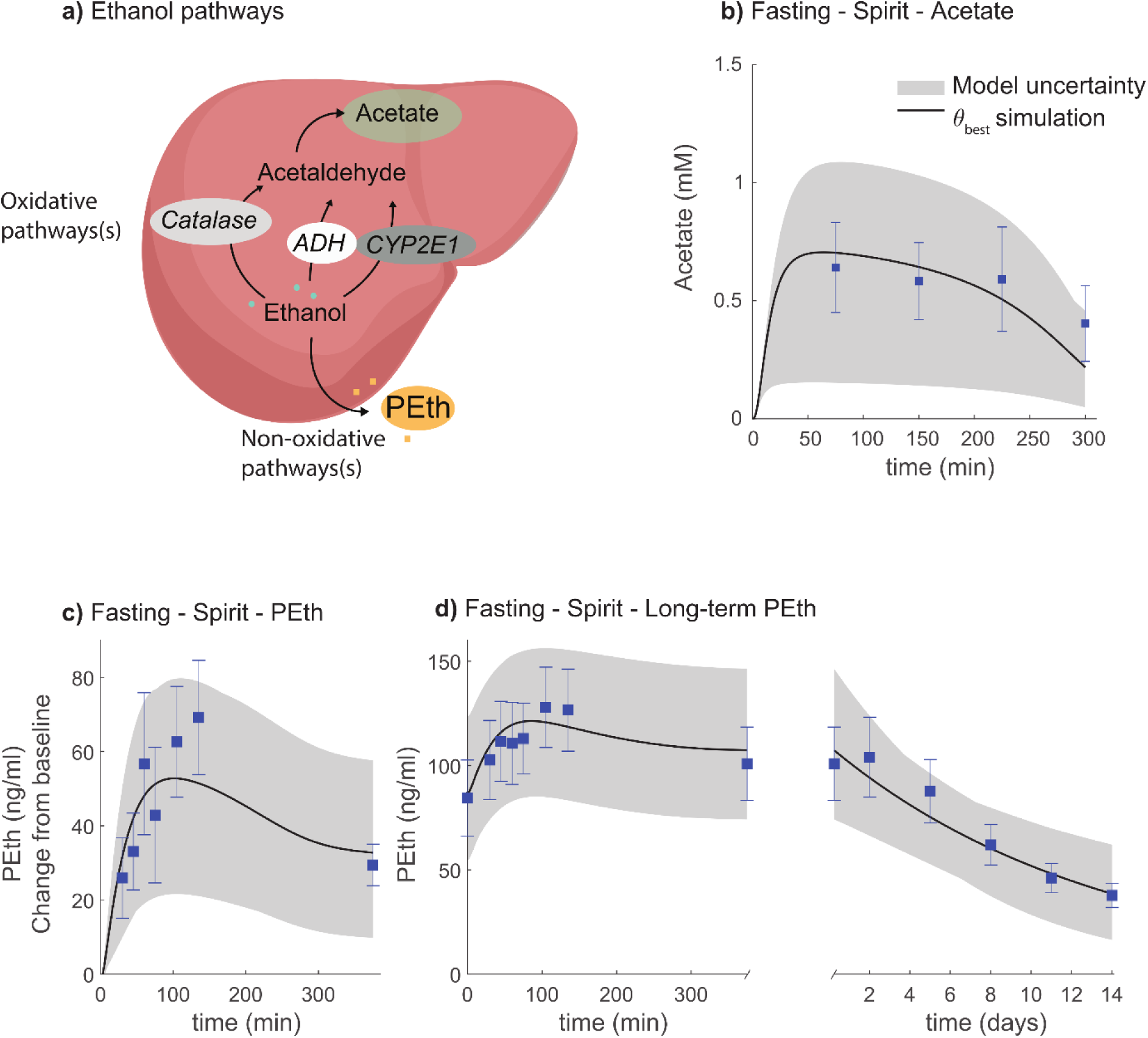
Model agreement to derivatives of oxidative and non-oxidative ethanol breakdown data for various interventions. The solid line is the best model fit, the shaded area is the model uncertainty, the blue boxes are the mean experimental data values with the error bars indicating the *standard error of the mean* (SEM). The alcohol was consumed orally through different beverages. **a)** Schematic of the oxidative and non-oxidative breakdown pathways. **b)** Estimated acetate from consumption of 0.28 L of 12.4 v/v % spirit blend (85 kcal) was consumed over 15 min. **c)** Estimated change of *phosphatidylethanol* (PEth), from the baseline value, from consumption of 0.71L of 6.5 v/v % spirit blend (247 kcal) over 15 min. **d)** Weighted mean behavior PEth from two groups consuming different beverages. Group 1 (n=16) consumed 0.72L of 3.25 v/v % spirit blend (251 kcal) over 15 min. Group 2 (n=11) consumed 0.71 L of 6.5 v/v % spirit blend (247 kcal) over 15 min. The combined PEth is the weighted mean behavior for both groups.

Firstly, for the short-term, the model agrees with data of PEth levels, presented as the change from the baseline value, directly after beverage consumption (Fig. 5c). Secondly, for the long-term, the model describes both the fast PEth plasma dynamics directly after beverage consumption (Fig. 5d, left), and subsequent slower long-term breakdown over several days (Fig. 5d, right). For the long-term dataset, Javors *et al*. presented the absolute PEth plasma concentration (ng/mL) for a group consisting of participants with different drinking schemes, low and high consumption, (Fig. 5d). As the low consumption group has more subjects (n=16), the combined group (n=27) is mostly influenced by the behavior of this group. The combined data is thereof a weighted mean behavior.

In summary, the model was capable of simultaneously describing data from different ethanol derivatives, in addition to the already shown agreement to gastric-emptying and plasma ethanol dynamics. In the following, the model is utilized to make predictions of scenarios not present in the estimation dataset.

### The model can predict independent validation data

The model was used to make predictions of the behavior of different attributes, and these predictions were validated against experimental data that was not used during model training (Fig. 6). The model predicts the gastric emptying rate following consumption of; 150 mL water (Fig. 6a), 150 mL glucose liquid containing 67 kcal (Fig. 6b), and 400 mL glucose liquid containing 200 kcal (Fig. 6c). These data were presented by Okabe *et al*. (29,50). As seen in the estimation data (Fig. 2), the total caloric content is the driving factor in decreasing the rate of gastric emptying. Predictions of plasma ethanol following consumption of an alcoholic beverage are compared with experimental data from Sarkola *et al*. (53) (Fig. 6d).

**Figure 6:**
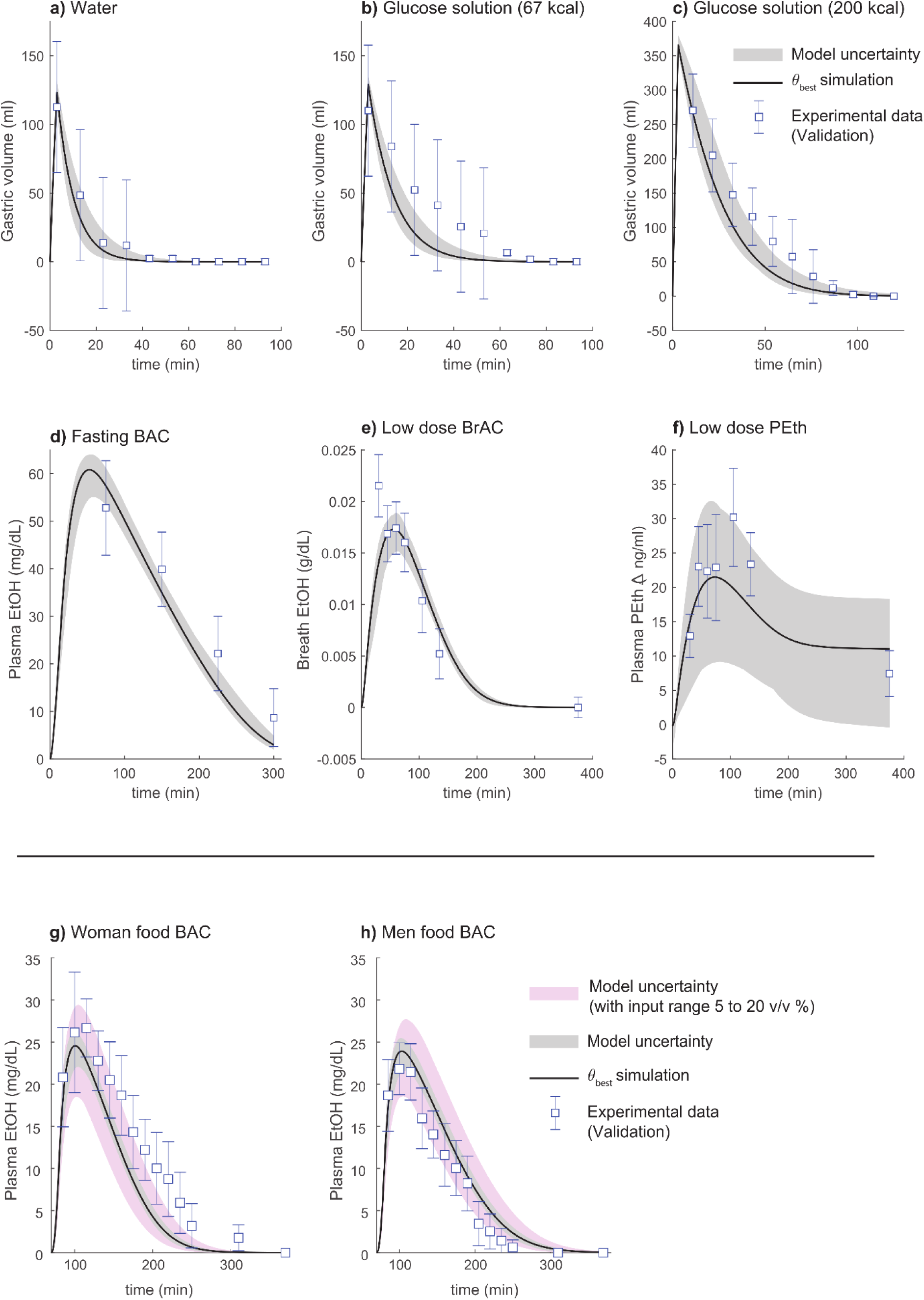
Model predictions compared to data for various interventions. The solid line is the best model fit, the shaded area is the model uncertainty, the white boxes are the mean experimental data values with the error bars indicating the *standard error of the mean* (SEM) – the white box indicates that the data was used for model validation. **a)** gastric emptying of 150 mL water, **b)** gastric emptying of 150 mL glucose liquid containing 67 kcal, **c)** gastric emptying of 400 mL glucose liquid containing 200 kcal, **d)** plasma ethanol levels from consumption of 0.48L of 10.0 v/v % spirit blend (149 kcal) over 15 min, **e)** breath ethanol levels from consumption of 0.72L of 3.25 v/v % spirit blend (251 kcal) over 15 min, **f)** change of PEth levels, from the baseline value, from consumption of 0.72L of 3.25 v/v % spirit blend (251 kcal) over 15 min, **g)** plasma ethanol levels in women after consumption of 0.20L of 12 v/v % spirit blend (36 kcal) over 10 min after eating a meal constituted of 555 kcal, **h)** plasma ethanol levels in men after consumption of 0.24L of 12 v/v % spirit blend (43 kcal) over 10 min after eating a meal constituted of 555 kcal. The pink uncertainty area in **g**-**h** was obtained by varying the drink concentration between 5-20 v/v%, as the details of the drink composition were not reported by Frezza et al. 1990.

The model could also predict data from a study by Javors *et al*. (52), of both BrAC (Fig. 6e) and the behavior of PEth levels as an increase from the baseline value (Fig. 6f), following consumption of an alcoholic beverage. Furthermore, the model could also predict sex-specific experimental data from Frezza *et al*. (54), of drinks consumed in combination with food for women (Fig. 6g) and men (Fig. 6h). One can observe that the model predicts a greater peak BAC amplitude for women. The participants in the Frezza *et al.* study were given the same dose of alcohol and the same size of the meal. Since the details of the drink composition were unreported, we show the model predictions using both a fixed drink concentration of 12.5 v/v% (Fig. 6g-h, grey area) and with different drink concentrations ranging from 5-20 v/v% (Fig. 6g-h, pink area).

### The model can be used to predict the effect of long-term alcohol use on PEth and differences in alcohol patterns

To highlight model usability, we used the validated model to highlight two different areas of use. The first example was to show how short-term drinking habits affect the long-term values of the alcohol consumption marker PEth (Fig. 7b, d). The second example highlights model personalization and how sex-specific anthropometric differences affect plasma ethanol dynamics in different drinking scenarios (Fig. 7a, c).

**Figure 7:**
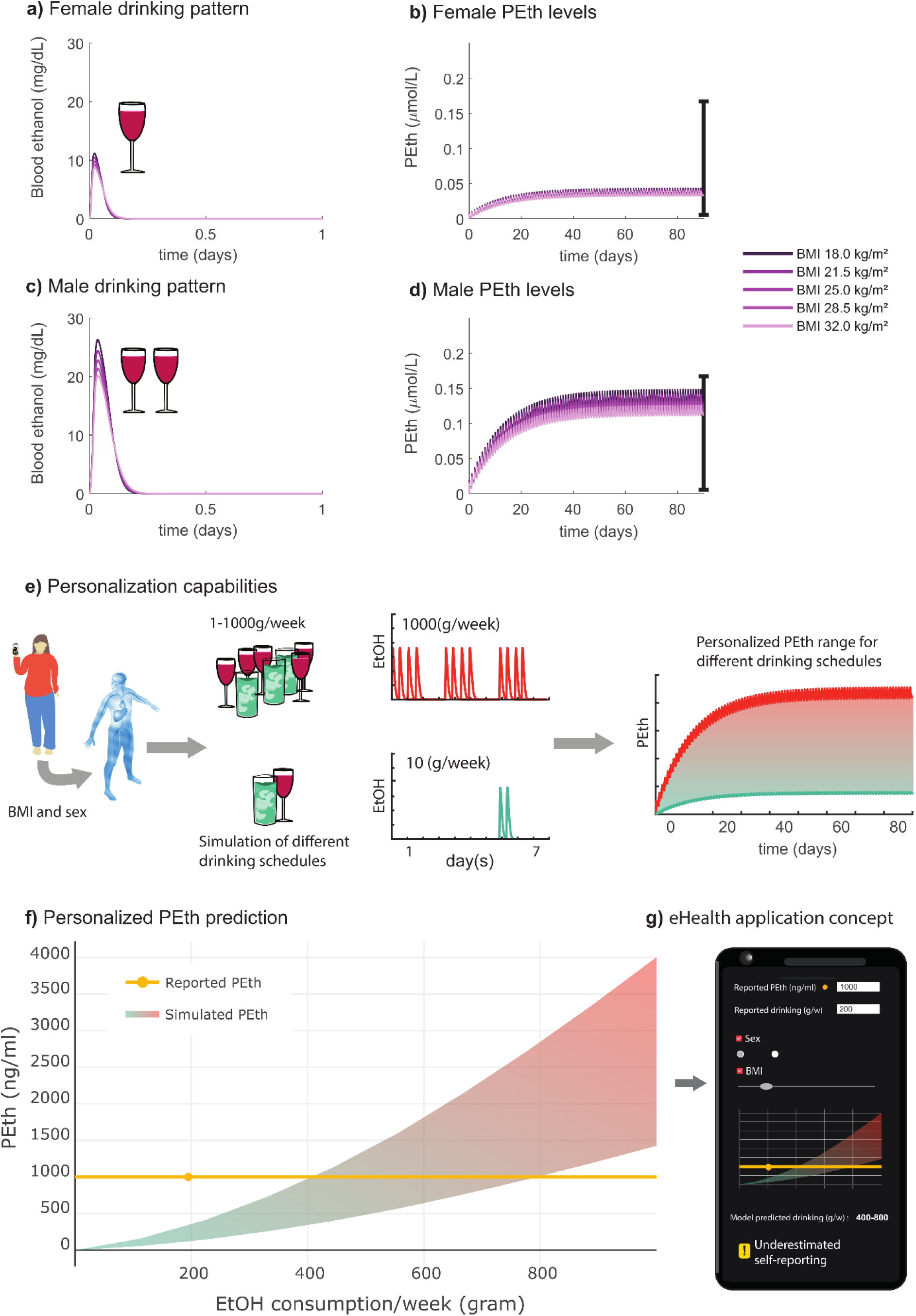
Model simulations of long-term plasma ethanol and PEth dynamics. *M*odel simulations of the short-term ethanol dynamics with corresponding long-term progression of PEth levels for two different drinking patterns, one for males and one for females, in the three month long drinking intervention presented in Kechagias *et al.* (24). A simulation approach for determining alcohol consumption is also shown. **a)** Model simulated ethanol dynamics during consumption of one standard glass of wine (15 cl, 13.5 v/v %) together with a meal consisting of 920 kcal by the female participants. The simulations were done for participants with different body mass indexes (BMIs) (18-32, light-to-dark purple lines). **b)** Model simulation of *phosphatidylethanol* (PEth) for females consuming one standard glass of wine every day for 90 days and for the five different body sizes, BMIs 18-28. The error bar represent the PEth range reported in Kechagias *et al*. (24). **c)** Model simulated ethanol dynamics during consumption of two standard glasses of wine (30 cl, 13.5 v/v %), in one sitting, together with a meal consisting of 1120 kcal, by the male participants. **d)** The corresponding model simulation of PEth progression for males consuming two standard glasses of wine every day for 90 days and for the five different body sizes, BMIs 18-32. The error bar represent the reported PEth range by Kechagias *et al*. (24). **e)** The simulation approach for determining alcohol consumption based on a given PEth level is shown. For a specific individual, defined via sex and BMI, the model also allows for simulations of different drinking patterns which can correspond to different amounts of alcohol consumed per week (1 to 1000 g/week). The weekly consumption could then be simulated for a longer time such that the steady-state PEth levels for the specific drinking schedule could be acquired. **f)** Using this approach for different amounts of consumption, the model could predict a trajectory for what weekly consumption corresponds to which PEth level range. The simulated trajectory is shown as the gradient area. The yellow line is an example of a predicted miss-match in self-reporting and model simulated alcohol consumption, for a specific PEth level. The yellow dot indicates the corresponding PEth levels (1000 ng/mL) of a self-reported consumption of 200 g/week ethanol. By comparing with the area, we can infer that the self-reporting was faulty, as such PEth levels require at least a consumption of ~400 grams/week. **g)** Here the future aim to integrate the physiologically-based digital twin into an eHealth application is shown, with possible use in clinics to motivate treatment compliance for patients with heavy drinking problems.

To investigate how well the model could assess the effect of long-term drinking on PEth levels, the model was used to simulate the PEth study presented by Kechagias *et al.* (24). In the study, 44 participants drank wine (13.5-14 v/v%) every evening (16.0-16.5 g ethanol, 1.3 standard drinks for women, and 32-33g ethanol, 2.7 standard drinks for men) for 3 months and their PEth levels were measured after 3 months. We simulated the study for both females and males with different anthropometric data (*body mass index* (BMI) 18, 22.5, 25, 28.5, and 32 kg/m^2^). This was done by varying the participant’s weight. For females, we simulated the consumption of one glass of wine per day together with a dinner containing 920 kcal (40% of recommended daily intake for a medium-active young adult female, in Sweden). This simulation was used to predict the daily plasma ethanol levels (Fig. 7a) as well as the long-term PEth levels after three months of daily drinking (Fig. 7b). For males, we simulated the consumption of two glasses of wine together with a dinner containing 1120 kcal (40% of recommended daily intake for a medium active young adult male, in Sweden). This simulation was used to predict the daily plasma ethanol levels (Fig. 7c) as well as the long-term PEth levels after three months of daily drinking (Fig. 7d). We compared the model-predicted long-term PEth values with the PEth range presented by Kechagias *et al*. (Fig. 7b, d). The model predicted PEth values for all subjects, with different anthropometrics, to be within the reported range of the Kechagias *et al*. measured values (0.007-0.17 *μmol*/*L*).

To illustrate the personalization capabilities of the model we have made simulations of either male (Fig. 7a) or female (Fig. 7c) anthropometrics, with varying BMI between 18 to 32 kg/m^2^. The consumption of one glass of wine, by females, yields a plasma ethanol level (11 mg/dL, BMI 18), which is less than half of that of males consuming 2 glasses of wine (27 mg/dL, BMI 18 kg/m^2^). This is explained by the relative higher ability of the meal (920 kcal for females and 1120 kcal for males) to lower the plasma ethanol for females. As males consume double the volume of wine the slightly bigger meal cannot slow down the appearance of ethanol in plasma as effectively as in the case of the females, resulting in a higher response. For both females and males, we observe a lower peak value with increasing body size (higher BMI value).

As proof of concept, we also showcase the model’s capability of predicting different weekly alcohol-consumption and the corresponding PEth levels (Fig. 7e, f). For a specific individual, defined via sex and BMI, the model can predict the long-term PEth levels for continuous drinking of various drinking schedules, corresponding to different amounts of weekly intake (g/week) (Fig. 7e). Using this approach, the model can predict a trajectory (gradient area) of what range of PEth levels correspond to which weekly consumption. Clinically, such a trajectory could be compared to a reported value (Fig. 7f, yellow dot), giving an indication if patients fail to accurately self-report consumption (Fig. 7f). In Fig. 7g we show as a concept, the further integration into an easy-to-use eHealth application, which we will create in the future.

In summary, our validated model can simulate long-term alcohol consumption and predict clinically relevant levels of PEth. Using the model, highly personalized predictions based on anthropometric data could also be predicted, showing the effect on different types of drinking habits. The model could thus be used in a wide variety of applications to assess the level of alcohol consumption e.g., to validate reported consumption via an AUDIT.

## Discussion

A mathematical model that can be personalized into a physiologically-based digital twin for alcohol consumption capable of simulating real-life drinking scenarios was developed. The model could simultaneously find agreement to: i) gastric emptying data for beverages with different volumes and caloric content (Fig. 2), ii) plasma ethanol in response to various drinking challenges (with and without food consumed) from different individuals (Fig. 4), and iii) data for markers of metabolized ethanol (Fig. 5).

Furthermore, we validated the model by making predictions of independent data consisting of gastric emptying, BAC, BrAC, and PEth (Fig. 6). Four different hypotheses of meal effects on plasma ethanol were also tested (Fig. 3), where only one hypothesis agreed with the experimental data (H1). The validated model was then used to investigate the effect of anthropometric differences on the short-term plasma ethanol dynamics (Fig. 7). The model predicts the long-term dynamics of PEth in response to daily consumption of alcohol (Fig. 7). These long-term predictions accurately describe the 90-day end of trail PEth range reported by Kechagias *et al*. (24) (Fig 7). Finally, we investigate model sensitivity subjected to the model parameters (Supplementary Materials “1 Parameter identifiability“) and the model inputs (Supplementary Materials “2 Analysis of the impact of the model inputs on ethanol dynamics”).

The model was trained and validated to a total of 10 different study datasets, which include different conditions, such as; different beverage volumes, ethanol content, food consumption or fasting, and sex-specific parameters (Fig. 2, 4-6) (24,28,29,33,37,50–54). An overview of the data is given in the Supplementary Materials, “S4 Usage of experimental data”. The model could explain all estimation and validation data for both ethanol and PEth to a satisfactory level (Fig. 4, 6). Furthermore, the model also describes the plasma acetate level following an alcoholic drink (Fig. 5b), with data from Sarkola *et al*. (53). Altogether, the model sufficiently describes the ethanol dynamics following consumption of beverages of different volumes, concentrations, time of consumption, in combination with food, and for individuals with different anthropometric data. With this model, we can make further simulation-based investigations of the impact of non-measured drinking series (see e.g. Supplementary Figure 22), to unravel the influence of non-alcoholic drinks on the BAC profile.

While the model passed a χ^2^-test for all estimation data, it is worth pointing out some aspects of the data that the model did not fully capture. Most notably, the model did not describe the observed amplitude in BAC data for some datasets. For instance, from the studies by Kechagias *et al*. and Jones *et al*., different BAC dynamics (Fig. 4d-e, 4g-h) were reported for the same consumption of 0.3 g ethanol/kg bodyweight for subjects with similar anthropometrics (33,37). As all the drinking challenges presented to the model are fitted with the same set of parameter values, the model is not able to account for differences in BAC behavior for similar drinking challenges (Fig. 4d-e). It was also observed that the model does not quite capture the dynamics of the peak in the data for beer and spirit consumption (Fig. 4a, c) from Mitchell *et al.* (51). This could be due to us estimating the caloric content incorrectly (see Supplementary Materials “3 Input estimations”), which in turn could make the gastric emptying rate of beer too fast and spirit too slow. Also, in the case of the beer data (Fig. 4a), the potential effect of carbonation on the rate of gastric emptying (58–60) could potentially influence gastric emptying to allow better plasma ethanol dynamics for the beer data.

In the validation data, “food-spirit woman” Fig. 6g, the model predicts that the BAC profile is too low compared with the experimental data. This is likely a result of the assumption of the drink ethanol concentration and the anthropometric data (see Supplementary Materials “3 Input estimations”). Anthropometric data such as BMI, and subsequent blood volume, impacts the ethanol dynamics (61) (Supplementary Figure 21). As a number of anthropometric data were not reported by Frezza et al. (54), we chose to use the anthropometrics of an average woman in Italy for the year 1990. Furthermore, no drink ethanol concentration was given, and we, therefore, assumed a drink concentration of 12.5 v/v% (Fig. 6g, grey area). These anthropometric values could be too large, resulting in a larger total blood volume, or the drink concentration could be different, which would then explain the low BAC profile. As these values could not be validated, we chose to stay consistent with the approach of estimating missing anthropometrics, even though these values might have been slightly incorrect. Instead, we varied the drink concentration between reasonable concentrations for alcoholic drinks (5-20 v/v%) since the concentration was not given (Fig. 6g, pink area). Given these alternative assumptions of the drink concentration, the model simulation can describe the experimental data.

Furthermore, individual, and sex-specific differences in the expression and phenotypes of ethanol metabolizing enzymes exist and could contribute to differences in ethanol plasma dynamics. The breakdown of ethanol is primarily enabled by enzymes such as *Alcohol dehydrogenases* (ADH), catalase, and *cytochrome P450 2E1* (CYP2E1). There are some sex-specific differences in the expression of hepatic ADH (62), also there are differences in ADH expression in individuals from different genetic backgrounds (63). These aspects were not included in the presented model but could be included in the future. By including this type of genetic and sex-specific differences, and other covariates, we could further improve the model adaptability when describing new cohorts.

Unlike previous models, our model has integrated gastric emptying dynamics to account for the influence of the caloric content, drink volume, and meal effect on gastric emptying (42,64–66). While newer models have been developed to include dynamics of the stomach, they are limited to only considering alcoholic drinks (43,45) and not the consumption of drinks paired with food. This makes them less usable when describing a real-life setting, where alcoholic drinks are often mixed with food and non-alcoholic drinks.

Several different hypotheses regarding the meal effect on gastric emptying were tested. The effect of meals on plasma ethanol dynamics has been discussed in various studies (33–36,55) and the literature is not unanimous on the underlying mechanism. Therefore, four different hypotheses (H1-H4) of the meal effect were introduced (Fig. 3a). In the first hypothesis (H1) food can encapsulate ethanol while present in the stomach (33). As the food is processed ethanol can be released back into the stomach and thus again become available for future absorption in the intestines. In the second hypothesis (H2) the contents of the stomach (including the liquid) are held longer in the stomach, due to the need for the food to be processed. In the third hypothesis (H3), H2 is extended to also include gastric ADH (33). In the final hypothesis (H4) the processing of the food recruits more blood to the gastric area which leads to both a faster uptake of alcohol and clearance due to increased activity of liver ADH (35,36,55). These hypotheses were tested by evaluating their agreement with the experimental data. It was found that only H1 had an acceptable agreement with data, and H2-H4 had to be rejected (Fig. 3b).

While the rejected hypotheses show that the corresponding mechanisms were not sufficient to explain the available experimental data, the mechanisms might still exist. For example, studies have shown that a meal reduces the amplitude and increases the clearance of plasma ethanol at least when ethanol is administrated intravenously (36,55). In this situation, additional mechanisms like the ones in H4 might be required. The reason is that consumed food will have limited physical interaction with alcohol in the stomach, likely meaning that the processing of the food recruits more blood to the liver. However, intravenous infusion of alcohol is not normal drinking, which the model is aimed at describing. Since these mechanisms were not needed for describing the estimation data, the mechanisms from H4 were not included in the final model. Another hypothesis we tested (H3) was the presence of gastric ADH, as there is a consensus that ADH is present in the stomach (35,67–69). In summary, these mechanisms were not able to describe the experimental data and they were therefore not included in the final model. The final model corresponds to H1 and can describe all the available experimental data sufficiently well.

In agreement with the work of Okabe *et al.* (28,29), the total beverage caloric amount was used as the driving factor for gastric emptying. This concept has been applied also in previous studies (32,70,71). This has been debated since other parameters are thought to be important such as; viscosity, osmolality, and nutrient composition (31,71,72). However, the effect of these factors appears to be outweighed by the total amount of non-ethanol calories, which was the only effect included in the model driving the liquid gastric emptying. Moreover, gastroparesis and the glucose control system have previously been identified as important factors for gastric emptying in diabetic patients (73,74).

Some simplifications of the gastric emptying rate of foodstuff were required. Firstly, the caloric density for the different macronutrients (carbohydrates, protein, and fat) was not differentiated. There are reported cases of macronutrients affecting the plasma ethanol level differently (33,34). Due to the small difference and macronutrient information not being consistently reported in studies, it was not included in the framework. Secondly, there are additional variables describing the meal that were not included in the model. One such variable is the viscosity of food products, although the caloric content has been shown to have a larger impact than the viscosity on gastric emptying (75). The caloric content was therefore chosen to be the driving factor.

Thirdly, another simplification is the potential ability of different food items to physically hinder ethanol absorption after consumption. The exact timing of the largest effect of food on the absorption is not known, which was reported by Franke *et al.* (76). Although the effect is reported, there is not enough data on the mechanisms to quantify the effect on the plasma ethanol rate of appearance. Lastly, an additional measure that could be considered is the fiber content of a meal. Including the fiber content in the model would allow for distinguishing between “liquid” and “solid” meals, and by extension the rate of gastric emptying of solids (77,78). However, it has been reported that the emptying rate of liquids remains mostly unaffected in the presence of foodstuff (30,79). The fibrous contents has previously not been reported to influence the ethanol rate of appearance in plasma, and it was not included here. In summary, there are a lot of reported mechanisms in literature, but they were not needed for the model to be able to explain data sufficiently well. Additional data would be needed to include these other mechanisms in our digital twin.

Simulation-based models, such as physiologically-based digital twins have not yet been explored in the context of eHealth applications aimed at reducing dangerous drinking habits. A variety of eHealth and smartphone applications have shown tentative promise as tools for reducing alcohol consumption (11,80–82). These applications mainly focus on either supporting, altering, or educating individuals about their drinking habits. More similar to the presented simulation-based model there are studies investigating eHealth applications and interventions integrating the use of eBAC calculators (18–21,83). In a study by Gajecki *et al.* (83), they did not observe an impact on drinking behavior using a smartphone application showing real-time assessment of eBAC, and simulations of predicted eBAC during a future drinking event was tested in a cohort of non-treatment-seeking university students. However, a later study by the same group Berman et al. (18), showed a positive effect using the same application in a group of hazardous drinkers from a similar type of cohort of university students. Furthermore, some studies have shown that the BAC calculator ‘apps’ commercially available, are unreliable, and there is a need for more evidence-based applications (20,21). One critical element we believe is required for BAC based applications to have an impact, is to connect short-term drinking to long-term consequences, something the physiologically-based digital twin is able to provide.

The validated model could predict PEth levels based on short-term consumption habits (Fig. 7b, d). The use of plasma PEth as a biomarker for alcohol consumption is widely used in clinical practice, sometimes as a validation to self-reporting systems (such as AUDIT) (84,85). As a proof-of-concept, simulations of long-term PEth levels in response to different drinking patterns and anthropometrics were preformed (Fig. 7 b, d), as well as a tool to evaluate self-reported weekly drinking in combination with measured PEth levels (Fig. 7 e-g).

In the future, a personalized model capable of predicting the long-term PEth trajectories given short-term drinking habits might prove useful for physicians working with patients, where distinguishing between moderate, and heavy drinking is important, e.g., liver-transplant patients. In the future, we would like to further validate the long-term PEth model, as well as expanding the ethanol degradation pathways in the model by adding alcohol consumption driven changes of CYP2E1 expression. With the intention to show how long-term exposure might lead to dysregulated metabolism and *alcohol related liver disease* (ALD).

There are mathematical models that describe different processes related to alcohol consumption, each with their strengths and limitations. The first kind is the gastric emptying models, which currently do not detail any interactions with the BAC profile following consumption of alcohol (38–40). On the other hand, the existing BAC models (41–45) fail to account for real-life drinking situations, (i) timing differences of gastric emptying, (ii) the influence of beverage caloric content, and (iii) meal effect on the BAC profile.

A second kind of already existing model type, are the minimal plasma ethanol models for determining kinetic parameters of ethanol dynamics (41,42). While these minimal plasma ethanol models can be used to estimate BAC, they are not useful when handling varying and combined ingestion of drinks and food. In contrast, there exist models of ethanol dynamics that include more detailed descriptions of the physiology (43,44). While these models describe the whole-body context of ethanol, they do not include a mechanistic description of gastric emptying and food consumption. A model by Moore *et al.* (45) includes the dynamics of ethanol absorption and the effect of a meal in non-human primates but it lacks a mechanistic description of gastric emptying, and can thus not explain the different gastric emptying rates of drinks with different caloric content. Furthermore, the model by Moore et al. does not describe long-term markers like PEth.

Lastly, there are also models connecting short-term ethanol dynamics to PEth. In a study by Simon (46), the Widmark model was extended to include a description of plasma PEth dynamics. While presenting a framework for long-term PEth estimation, this work is unable to account for the real-life drinking scenarios previously listed.

Future versions of our model could include disease progression or long-term regulatory changes of hepatic enzymes. There already exist mathematical models describing the expression of CYP2E1 (86), but not in the context of increased alcohol consumption. Moving beyond ODE models there exist other approaches. *Artificial intelligence* (AI) and *machine learning* (ML) solutions are powerful methodologies to extract useful information from data, which could be applicable in the context of alcohol consumption. There exist studies using AI approaches in the context of detecting harmful alcohol use (87). While powerful, these tools require a critical amount of data. Furthermore, the ML/AI models are black-box models, in contrast to the mechanistic ODE models which are based on mechanistic hypotheses. To train the ODE models, less data is necessary when compared to when training ML/AI models. Furthermore, since the parameters in the ODE models often have a direct biological interpretation, the models are easier to personalize.

There are some model limitations and delimitations that should be discussed. First, we consider limitations in the data, and then limitations in the model. All data used in this project was acquired via digitization (extracted from figures in peer-reviewed publications), and thus a negligible difference from the actual data might exist. However, the qualitative information in the dataset is the same. In addition to this, some of the studies did not report anthropometrical data of the subjects or details regarding several aspects of the experimental setup e.g., reported drinking time. Also, for some datasets mean values were calculated in a mixed cohort including both female and male participants (52,53). To take this into consideration we opted to estimate the mean blood volume based on the sex distribution. Also, for some studies the caloric content was not reported, leading to some assumptions regarding the consumed beverage. The caloric content is of importance as the gastric emptying module is the major contributor to the ethanol rate of appearance in plasma. For details regarding assumptions see Supplementary Materials “3 Input estimation”. A delimitation for the model is the oxidative breakdown of ethanol, where we currently only include data for acetate dynamics. Future versions of the model could extend this by e.g., also including acetaldehyde. Finally, the last limitation is that our digital twin does not have real-time data integration via wearable sensors. There are different types of digital twins, and only some of them require real-time data integration (47–49). We have not implemented such real-time integrations. Thus, the study presents an ‘offline’ digital-twin, or the offline portion of the calibration of a digital twin.

We herein present a physiologically-based digital twin, including a gastric emptying module, capable of describing plasma ethanol dynamics for a wide variety of drinking scenarios. The twin also connects short-term alcohol consumption to the long-term progression of the clinical marker PEth. We believe that the digital twin, with further validation and integration of more personalization (such as genetic variations in enzyme expression), could serve as a valuable clinical tool. For example, to test reported alcohol consumption in an AUDIT questionnaire, by evaluating PEth values in a personalized manner. Furthermore, this digital twin could function as a first simulation-based eHealth application of real-life drinking scenarios, with the goal of educating people about their alcohol drinking habits and their long-term effects.

## Methods

### Model description

Within this section, all the equations in the model are detailed. The final model is shown in Fig. 1c, and the equations are given in Equations (2) to (8).

Before detailing the equations, an example of an *ordinary differential equation* (ODE) is described. A typical ODE used in this work looks similar to Eq. (1).

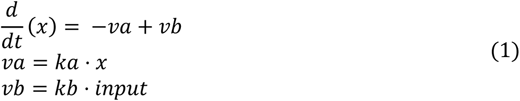

Here, *x* is a state in the model, *va* and *vb* are reaction rates, *ka* and *kb* are rate-determining parameters, and *input* is some input to the state. In other words, the amount of the state *x* is decreased by the reaction *va* with the speed *ka* and increased by the reaction *vb* with the speed *kb* depending on some input *input*.

#### The gastric emptying dynamics

The model describes the gastric emptying dynamics using the volume and the caloric content (Eqs. 2–3). While drinking, the variable *volDrinkPerTime* increases the volume (*Vol_Stomach_*) and the caloric content (*Kcal_Liquid_*) in the stomach. The liquid in the stomach is then emptied using the reaction *r*2. As the gastric emptying is driven by the total amount of (liquid) kcal the *Kcal_Liquid_* state is emptied over time. As the volume of the stomach can’t be equal to zero (zero division), *Vol_change_* is used to reach a zero value. Here, *SS_vol_* is a variable that defines the minimal volume of the stomach, set to 0.001 *dL*. The kcal consumed via food is set via the initial condition to the state *Kcal_Solid_* if a meal is consumed. The meal is emptied from the stomach using *rKcal_Solid_*. This reaction was originally presented by Tougas *et al.* (56), as a time-dependent expression, see Eq. (4). To make it compatible with the ODE format, we derived the derivative of the expression and scaled with the total consumed kcal to express the retention of kcal instead of *%*-retention, see *rKcal_Solid_* in Eq. (3). As the emptying is dependent on the total amount of solid kcal, *maxKcal_Solid_* keeps track of the initial (maximal) amount of meal kcal. On consecutive meals, the consumed kcals are added to *maxKcal_Solid_*, and *Kcal_Solid_*. Furthermore, *timeElapsed* keeps track of the time passed since the last meal (in minutes). Finally, *kcalSolid_vol_* transforms the kcal into a volume using the average caloric energy density of 4 kcal/g and assumes a density of 1 *g/ml* (dividing with 100 to go to *dL*).

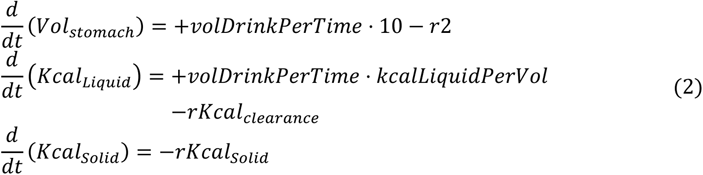

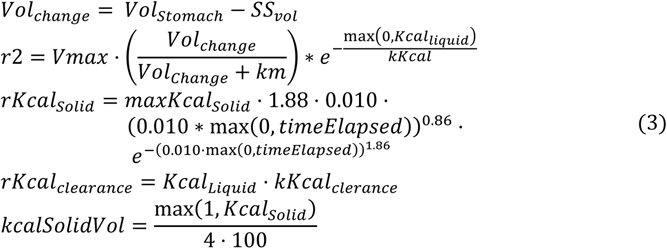

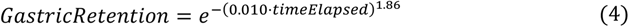

#### Ethanol dynamics

In the stomach, ethanol can interact with consumed meals. *EtOH_Pool_* allows for concentration-driven encapsulation of ethanol into the food (*rPool_In_* binds and *rPool_Out_* releases ethanol), effectively making it unavailable for the stomach while encapsulated. As the *Kcal_Solid_* is digested the volume of the pool decreases and ethanol is released back into the stomach again. When the kcals from food (*Kcal_Solid_*) have been fully digested (i.e. reached zero), *maxKcal_Solid_* and *EtOH_Pool_ were also set* to zero. With the consumption of alcoholic beverages (EtOHConc *>* 0), the concentration of ethanol in the stomach *ConcEtOH_Stomach_* is increased via *rDrink_EtOH_*. Here, *concDrink* converts *EtOHConc* (*v/v%*) to *mg/dL*, using the conversion factor 789.1 *mg/dL*. The ethanol can be encapsulated (*rPool_In_*) or released from the encapsulation (*rPool_Out_*). As the liquid is leaving the stomach (*r*2) the ethanol mass is introduced to the intestines *MassEtOH_Intestines_*. From the intestines, ethanol can be absorbed into the blood/plasma *BloodConc* via *r*3 and be eliminated from the body (*r*4). In the blood/plasma compartment, the ethanol can be eliminated via enzymatic activity in the liver *r*5. The model includes the ADH (*vADH*) and CYP2E1 (*vCYP*2*E*1) pathways. As ethanol is treated as a concentration, there is a volume scaling between the blood volume (*V_Blood_*) and the liver volume (*V_Liver_*).

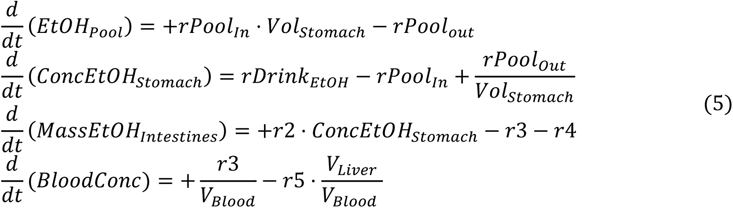

Here, *rPool_In_* and *rPool_Out_* control the diffusion of ethanol into the food pool. This diffusion can only occur if there is food in the stomach. Thus, if there is no food (if *Kcal_Solid_ <*= 1) or the total volume of the stomach is small (*Vol_Stomach_ <* 2·*SS_Vol_*), *rPool_In_* and *rPool_Out_ are set* to zero. The equations for *rPool_In_* and *rPool_Out_*, if there is both food and sufficient total volume in the stomach, are given in Eq. (6).

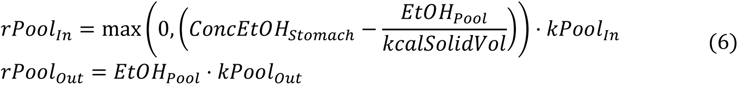

The other reaction rates used in the ODEs in Eq. (5) are described in Eq. (7).

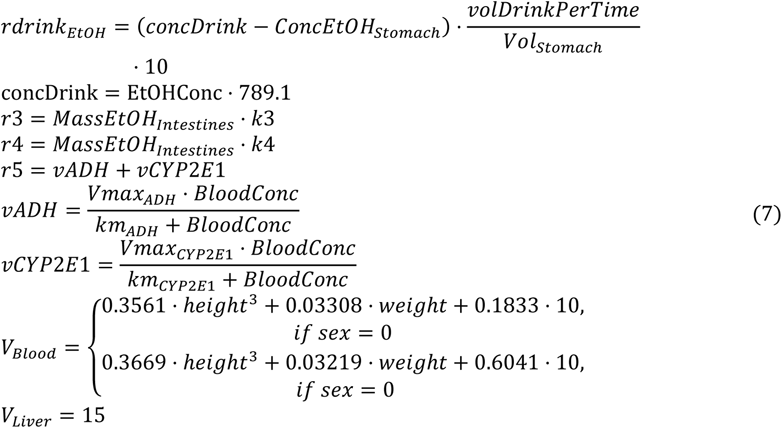

#### Ethanol metabolites

As ethanol is metabolized by ADH and CYP2E1 it is creating acetate *Plasma_Acetate_* via *r*5. Acetate is in turn cleared with *r*6. While present in plasma, ethanol can also be converted to *PEth* via *rPEth*. PEth is cleared from the plasma using *rPEth_clearance_*. Additionally, the PEth can be bound to lipids in the body and be temporarily stored. PEth is bound via *rPEth_bound_* and released via a concentration gradient *rPEth_release_*.

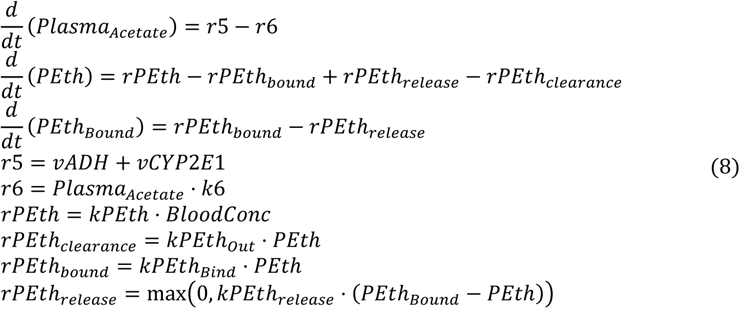

#### Initial values

It was assumed, in the model, that the person has no residual alcohol in the system. Furthermore, it was assumed that the model starts in a fasted state, with no kcal in the system, and that the residual volume in the stomach was 0.001 *dL*. However, the basal levels of PEth was allowed to be within a range for the experiments containing PEth, because Javors et al. reported levels of PEth beyond 0 (52). In this case, the initial values for the states PEth and PEth bound need to be calculated from the estimated basal PEth levels and the parameter values, see equation Eq. (8). Otherwise, the initial values of PEth and PEth_Bound_ were assumed to be 0. The initial values used are given in Table 1.

**Table 1:** Initial values.

| State | Initial values |
| --- | --- |
| $Vol_{Stomach}$ | $1.0000 \cdot 10^{-3}$ |
| $Kcal_{Liquid}$ | 0.0000 |
| $MaxKcal_{Solid}$ | 0.0000 |
| $Kcal_{Solid}$ | 0.0000 |
| $timeElapsed$ | 0.0000 |
| $EtOH_{Pool}$ | 0.0000 |
| $ConcEtOH_{Stomach}$ | 0.0000 |
| $MassEtOH_{Intestines}$ | 0.0000 |
| $BloodConc$ | 0.0000 |
| $Plasma_{Acetate}$ | 0.0000 |
| $PEth$ | *estimated* |
| $PEth_{Bound}$ | *estimated* |

#### Parameter values

This section gives the optimal parameter values for the connected model when estimated to the estimation dataset (columns *θ_est_*\*). Furthermore, the bounds used in the optimization for all parameters are also given (columns lower bound and upper bound), see Table 2. *km_ADH_* and *km_CYP_*_2*E*1_ were given bounds reported in literature (88). Lastly, the minimum and maximum parameter values for the model uncertainty are given (columns CI lower bound and CI upper bound).

**Table 2:** Parameter values and parameter estimation bounds.

| Parameter | $\theta_{est}^*$ | lower bound | upper bound | CI lower bound | CI upper bound |
| --- | --- | --- | --- | --- | --- |
| $kPEth$ | $1.4599 \cdot 10^4$ | $1 \cdot 10^{-5}$ | $1 \cdot 10^5$ | $6.3284 \cdot 10^3$ | $1.5557 \cdot 10^4$ |
| $kPEth_{out}$ | $1.5695 \cdot 10^2$ | $1 \cdot 10^{-5}$ | $1 \cdot 10^5$ | $6.0869 \cdot 10^1$ | $2.5765 \cdot 10^2$ |
| $kPEth_{bind}$ | $1.7416 \cdot 10^4$ | $1 \cdot 10^{-5}$ | $1 \cdot 10^5$ | $8.8872 \cdot 10^3$ | $2.3679 \cdot 10^4$ |
| $kPEth_{release}$ | $5.8081 \cdot 10^{-3}$ | $1 \cdot 10^{-5}$ | $1 \cdot 10^5$ | $9.7018 \cdot 10^{-4}$ | $1.1545 \cdot 10^{-2}$ |
| $kPool_{In}$ | $1.0155 \cdot 10^1$ | $1 \cdot 10^{-5}$ | $1 \cdot 10^5$ | $4.8248 \cdot 10^{-1}$ | $2.8715 \cdot 10^3$ |
| $kPool_{Out}$ | $4.0227 \cdot 10^{-1}$ | $1 \cdot 10^{-5}$ | $1 \cdot 10^5$ | $2.0875 \cdot 10^{-3}$ | $7.4358 \cdot 10^{-1}$ |
| $Vmax$ | $2.1545 \cdot 10^3$ | $1 \cdot 10^{-5}$ | $1 \cdot 10^5$ | $1.2948 \cdot 10^3$ | $2.4439 \cdot 10^3$ |
| $km$ | $1.7793 \cdot 10^4$ | $1 \cdot 10^{-5}$ | $1 \cdot 10^5$ | $1.6709 \cdot 10^4$ | $2.1276 \cdot 10^4$ |
| $kKcal$ | $1.5843 \cdot 10^2$ | $1 \cdot 10^{-5}$ | $1 \cdot 10^5$ | $1.1600 \cdot 10^2$ | $2.4235 \cdot 10^2$ |
| $k3$ | $8.4172 \cdot 10^1$ | $1 \cdot 10^{-5}$ | $1 \cdot 10^5$ | $5.4902 \cdot 10^1$ | $8.4172 \cdot 10^1$ |
| $k4$ | $8.3001 \cdot 10^2$ | $1 \cdot 10^{-5}$ | $1 \cdot 10^5$ | $7.1554 \cdot 10^2$ | $9.5080 \cdot 10^2$ |
| $k6$ | $1.3079 \cdot 10^{-1}$ | $1 \cdot 10^{-5}$ | $1 \cdot 10^5$ | $6.4762 \cdot 10^{-2}$ | $1.0000 \cdot 10^{20}$ |
| $Vmax_{ADH}$ | $9.6381 \cdot 10^{-1}$ | $1 \cdot 10^{-5}$ | $1 \cdot 10^5$ | $7.4245 \cdot 10^{-1}$ | $9.6381 \cdot 10^{-1}$ |
| $Vmax_{CYP2E1}$ | $1.7148 \cdot 10^{-1}$ | $1 \cdot 10^{-5}$ | $1 \cdot 10^5$ | $2.4640 \cdot 10^{-3}$ | $1.7366 \cdot 10^{-1}$ |
| $km_{ADH}$ | $9.2200 \cdot 10^0$ | $9.22 \cdot 10^{-1}$ | $9.22 \cdot 10^0$ | $6.7451 \cdot 10^0$ | $1.0220 \cdot 10^1$ |
| $km_{CYP2E1}$ | $3.6880 \cdot 10^1$ | $3.688 \cdot 10^1$ | $4.61 \cdot 10^1$ | $1.2489 \cdot 10^0$ | $3.7118 \cdot 10^1$ |
| $kKcal_{clearance}$ | $6.1804 \cdot 10^{-3}$ | $1 \cdot 10^{-5}$ | $1 \cdot 10^5$ | $1.0000 \cdot 10^{-20}$ | $1.4981 \cdot 10^{-2}$ |

#### Model inputs

This section lists the input values the model needs, see Table 3. A detailed overview of all the inputs provided to the model for each dataset is provided in the Supplementary Materials, see “3 Input estimations”.

**Table 3:** Input information to the model.

| Input variable | Description |
| --- | --- |
| $EtOHConc$ | Ethanol concentration |
| *volDrinkPerTime* | Consumed volume per minute |
| *kcalLiquidPerVol* | Amount kcal per liter beverage |
| *sex* | Male (1) or Female (0) |
| *weight* | Weight in kg |
| *height* | Height in meter |

#### Model outputs

This section lists the model outputs and the scaling performed. *yAcetate* is divided with 10.2 convert the concentration unit *mg/dL* to *mM*. *yBrAC* rescales the plasma concentration of ethanol *Blood_Conc_* into breath concentration using a linear correlation observed between BrAC and BAC measurements by Skaggs et al. (89). The additional division by 1000 is to go from *g* to *mg*.

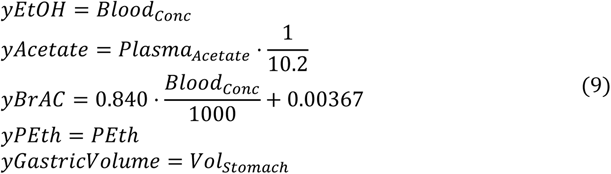

#### Modelling process

This model is the result of the iterative modelling cycle depicted in Fig. 1b. The model, the ODEs given above, aims to describe data and prior knowledge available, to be used as a reliable analytical tool of the system. The process of achieving such a model is tightly linked to the available experimental data, where data is used to evaluate and validate the model’s behavior. In this work, previously published data was used to describe gastric emptying (liquid and food), the rate of appearance of ethanol in the blood following consumption of beverages and food, and the rate of appearance of PEth following different drinking schemes. These data were divided into an estimation group, see Supplementary table 26, which was used to train and evaluate the model, see Fig. 2-5, and a validation group, see Supplementary table 27, which was used to validate the model see Fig. 6-7.

Using the evaluation data, the model behavior was iteratively evaluated and with each iteration, the model structure was adjusted against new data and literature knowledge. An example of this is given in Fig. 3, where 4 different hypotheses (H1-H4) of the food and ethanol’s rate of appearance in the blood were tested. After some iterations and the inclusion of more experimental data, H2-H4 could be rejected due to insufficiently describing the estimation data. Eventually, the model can satisfyingly showcase the tested properties, given the evaluation dataset. Thereafter, the model is evaluated against the validation data, which does not influence the model behavior during the evaluation phase. A reliable model should be able to describe new data of similar scenarios from new cohorts. A validated model can be used to make predictions of untested, or unmeasurable, experimental designs. Such a prediction is given in Fig. 7b and Fig. 7d, where the model is used to evaluate the PEth profile following daily consumption of wine, and the intermediate PEth profile from days 1-89 is predicted. As this prediction cannot be validated, unless a new experiment is carried out, the strength of the prediction is dependent on the evaluation and validation phase of the model design.

### Parameter estimation

All model analysis and simulation were performed in both MATLAB 2022b using the system biology toolbox (90). For model parameter estimation the *extended scatter search* (ESS) algorithm implemented in the *MEtaheuristics for systems biology and bIoinformatics Global Optimization* (MEIGO) toolbox was used (91).

Parameter estimation was done by quantifying the model performance, using the model output ŷ to calculate the traditional weighted least squares cost function defined as

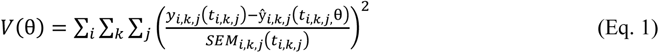

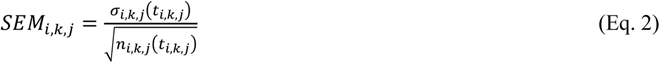

where, θ is the model parameters; *y_i,k,j_*(*t_i,k,j_*) is the measured data from a study *i*, and from on type of measure *k*, at time point *j*; ŷ*_i,k,j_*(*t_i,k,j_,* θ) is the simulation value for a given experiment setup *i,* type of measure *k,* and time point *j*. SEM is the standard error of the mean, which is the sample standard deviation, σ*_i,k,j_*(*t_i,k,j_*) divided with the square root of the number of repeats, *n_i,k,j_*(*t_i,k,j_*) at each time point. The value of the cost function, *V*(θ), is then minimized by tuning the values of the parameters, typically referred to as parameter estimation.

To evaluate the new model, a *χ*^2^-test for the size of the residuals, with the null hypothesis that the experimental data have been generated by the model, and that the experimental noise is additive and normally distributed was performed. In practice, the cost function value was compared to a *χ*^2^ test statistic, 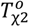. The test statistic *χ*^2^ cumulative density function,

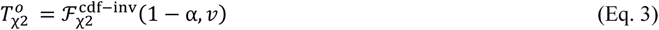

where 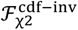 is the inverse density function; and α is the significance level (α = 0.05, was used) and *v* is the degrees of freedom, which was equal to the number of data points in the estimation dataset (193 in total, all timepoints over all experiments). In practice, the model is rejected if the model cost is larger than the *χ*^2^-threshold (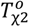).

### Uncertainty estimation

The model simulation uncertainty was gathered as proposed in (92) and is given in the Supplementary Materials, see “1 Parameter identifiability”. The model uncertainty is estimated by dividing the problem into multiple optimization problems, with one problem per model property (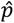). In this work, the property 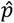 corresponds to either a simulation at a specific time point *j*, *ŷ*(*t_j,_*θ), or a parameter value 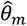. Each problem is solved by maximizing and minimizing the property value, while satisfying that the cost (*V*(θ)) is below the *χ*^2^-threshold (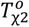). By identifying the maximal and minimal value of the model property (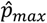 and 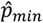), the boundary values of the property uncertainty area are found. Mathematically, this operation for the parameter values is formulated as,

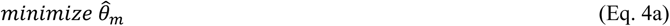

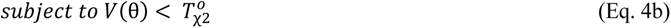

where 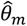 is minimized to find the lower value of the parameter, while also satisfying that the cost (*V*(θ)) is below the *χ*^2^-threshold (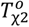). To find the upper bound of the uncertainty area the problem is maximized instead. In practice, the constraint (Eq 4b) can be relaxed into the objective function as a L1 penalty term with an offset if 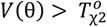.

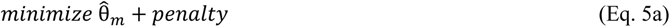

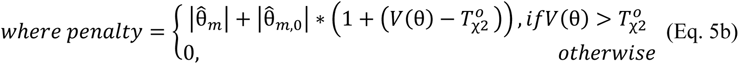

Here, the penalty is scaled with the initial value of the parameter, 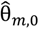 and the offset between the cost and the *χ*^2^-threshold (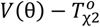). To maximize the parameter 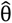, and thus finding the upper bound of the uncertainty area, the problem is solved as a minimization problem. This is done by substituting 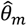 with 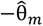 in the objective function. To solve the problem for the model simulation at a specific time point, *ŷ*(*t_j_,* θ), the problem is formulated as follows,

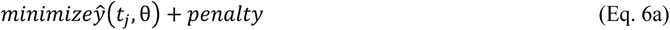

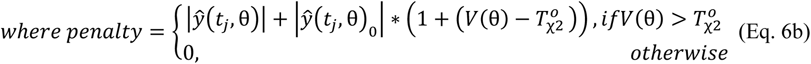

Here, the penalty is scaled with the initial value of the parameter, ŷ(*t_j_,* θ)_0_, and the offset between the cost and the *χ*^2^-threshold (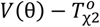). To maximize the model simulation at time point *j*, (*ŷ*(*t_j_,* θ), and thus finding the upper bound of the uncertainty area, the problem is solved as a minimization problem. This is done by substituting *ŷ*(*t_j_,* θ) with −*ŷ*(*t_j_, θ*) in the objective function.

### The experimental data used for the modelling

In this work, experimental data was collected from a wide variety of different published studies and used for model estimation and validation. This section will detail information about the studies included. Additionally, this information is condensed into two tables, one for estimation data and one for validation data, in the Supplementary Materials “3 Usage of experimental data”.

#### Gastric emptying

Three studies were included concerning the gastric emptying of liquids, all being works of Okabe *et al*. The first study describes the emptying rate of 500 ml liquids containing different amount of calories (28). The second study investigated how the total volume affects the emptying of isocaloric beverages (29). The third study investigates how calories from ethanol compare to non-alcoholic calories in isometric volume beverages.

#### Blood alcohol concentration

A variety of studies observing the BAC levels were included, some including the impact of a meal in combination with an alcoholic beverage. Mitchell *et al.* investigated the effect of consuming different alcoholic beverages with the same total alcoholic content (51). Sarkola *et al.* investigated an additional variant of alcoholic beverage (53). Kechagias *et al.* and Jones *et al.* both compared the BAC levels after consumption of an alcoholic beverage, with and without having a meal (33,37). Lastly, the work of Frezza *et al.* presents sex-specific differences in the BAC levels after consumption of an alcoholic beverage, where the ethanol contents were determined by the subjects’ body weight, in combination with a meal (54).

#### Blood alcohol derivates

Several studies measuring either acetate or PEth after ingestion of alcohol were included. Firstly, in a study by Sarkola *et al.* the appearance of blood acetate after consumption of a single alcoholic drink (53) was reported. Secondly, the time series of BrAC and PEth were published by Javors *et al*. for two drinks with different alcoholic content (52). Javors *et al.* also presented PEth measurements over a two-week period. Lastly, Kechagias *et al.* presented a study where the PEth level were measured before and after a 30-day period of daily consumption of wine (24).

## Data availability statement

All data used for model estimation and validation can be accessed from the original publications. We provide all model related data files and parameter values in our public code repository, DOI: 10.5281/zenodo.10104891.

## Code availability statement

All related scripts and datafiles are provided in our GitHub repository https://github.com/willov/ameta with a permanent copy available at Zenodo (DOI: 10.5281/zenodo.10104890). Additionally, a user interface for the model, through a Streamlit application, is provided. This application is available from our GitHub repository https://github.com/willov/alcohol_app with a permanent copy available at Zenodo (DOI: 10.5281/zenodo.10054299).

## Abbreviations

ADH: Alcohol dehydrogenase
AUD: Alcohol use disorder
AUDIT: Alcohol use disorders identification test
ALD: Alcohol related liver disease
AI: Artificial intelligence
BAC: Blood alcohol concentration
BMI: Body mass index
BrAC: Breath alcohol concentration
CYP2E1: Cytochrome P450 2E1
eBAC: Estimated blood alcohol concentration
ESS: Extended scatter search
HCC: Hepatocellular carcinoma
ML: Machine learning
MEIGO: MEtaheuristics for systems biology and bIoinformatics Global Optimization
ODE: Ordinary differential equation
PEth: Phosphatidylethanol
SEM: Standard error of the mean

## Acknowledgements

The computations were enabled by resources provided by the National Supercomputer Centre (NSC), funded by Linköping University.

The authors acknowledge financial support from: ALF Grants, Region Östergötland (PN, ME), Knut and Alice Wallenberg Foundation and Wallenberg Center for Molecular Medicine, Linköping University (PN), The Swedish Society of Medicine (PN), Bengt Ihre Foundation (PN), Magtarmfonden, Sweden (PN). GC acknowledges support from the Swedish Research Council (2018-05418, 2018-03319), CENIIT (15.09), the Swedish Foundation for Strategic Research (ITM17-0245), SciLifeLab National COVID-19 Research Program financed by the Knut and Alice Wallenberg Foundation (2020.0182), the H2020 project PRECISE4Q (777107), the Swedish Fund for Research without Animal Experiments (F2019-0010), ELLIIT (2020-A12), VINNOVA (VisualSweden, 2020-04711), and the Horizon Europe project STRATIF-AI (101080875). GC and WL acknowledge scientific support from the Exploring Inflammation in Health and Disease (X-HiDE) Consortium, which is a strategic research profile at Örebro University funded by the Knowledge Foundation (20200017). WL acknowledges support from the Area of Strength e-Health at Linköping University and Region Östergötland

The funders had no role in study design, data collection and analysis, decision to publish, or preparation of the manuscript.

## Author Contributions

**Conceptualization:** HP, CS, PN, ME, SK, PL, WL, and GC. **Formal model analysis:** HP, CS, WL, and GC. **Data curation:** HP, CS, and WL **Visualization:** HP, CS, WL, and GC. **Supervision:** PN, ME, SK, PL, WL, and GC. **Funding acquisition:** PL, WL, and GC.

All authors were involved in writing and reviewing the manuscript. All authors read and approved the final manuscript.

