## Supplementary Information for "A physiologically-based digital twin for alcohol consumption – predicting real-life drinking responses and long-term plasma PEth"

March 13, 2024

**4 Usage of experimental data 51**

### 1 Parameter identifiability

We tested the identifiability of the model parameters by formalizing an optimization (min or max) problem that would find the bounds of the parameter values, while also requiring the overall agreement to the data to be sufficiently good. During the optimization, we allowed all parameters to vary between the bounds defined in table 2, except for the parameter being tested which was allowed an unlimited range. We ran the optimization multiple times for all the parameter values at the Swedish national supercomputer center in parallel. We gathered the found optima and sorted them into a waterfall plot for all parameters (Supplementary Figures 1 to 19).

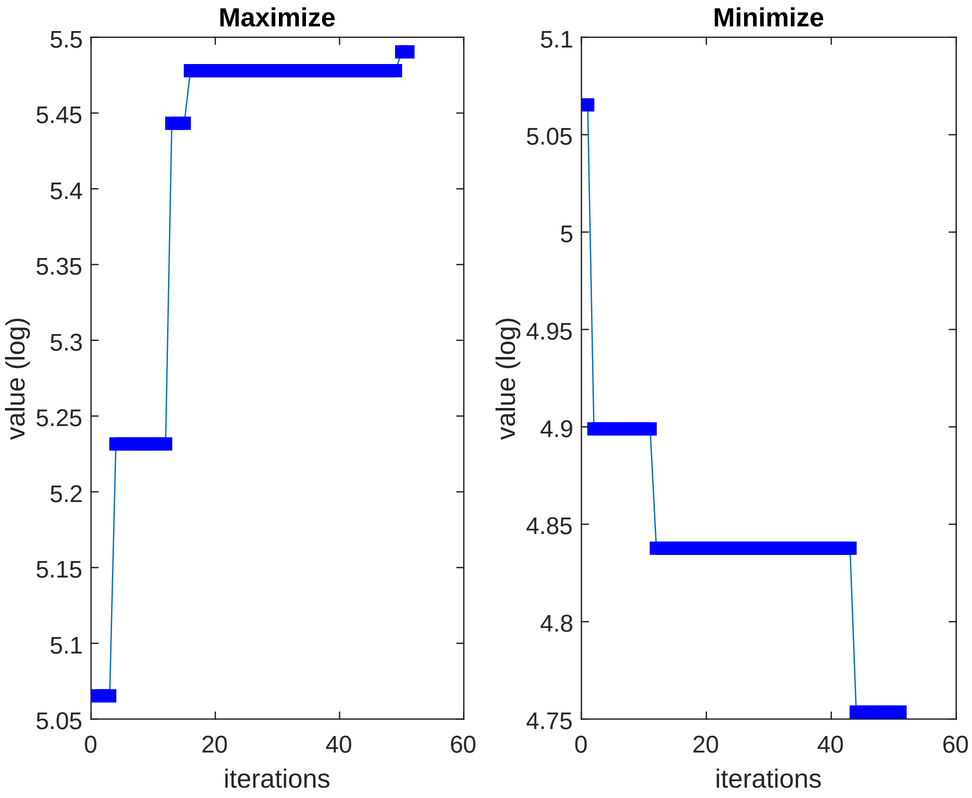

Supplementary Figure 1: The waterfall plot for the parameter k_kcal. The parameter value of k_kcal was both minimized and maximized to get the bounds, while still requiring the overall agreement to data to be sufficiently good (determined with a $\chi^2$-test). Parameter k_kcal is identifiable.

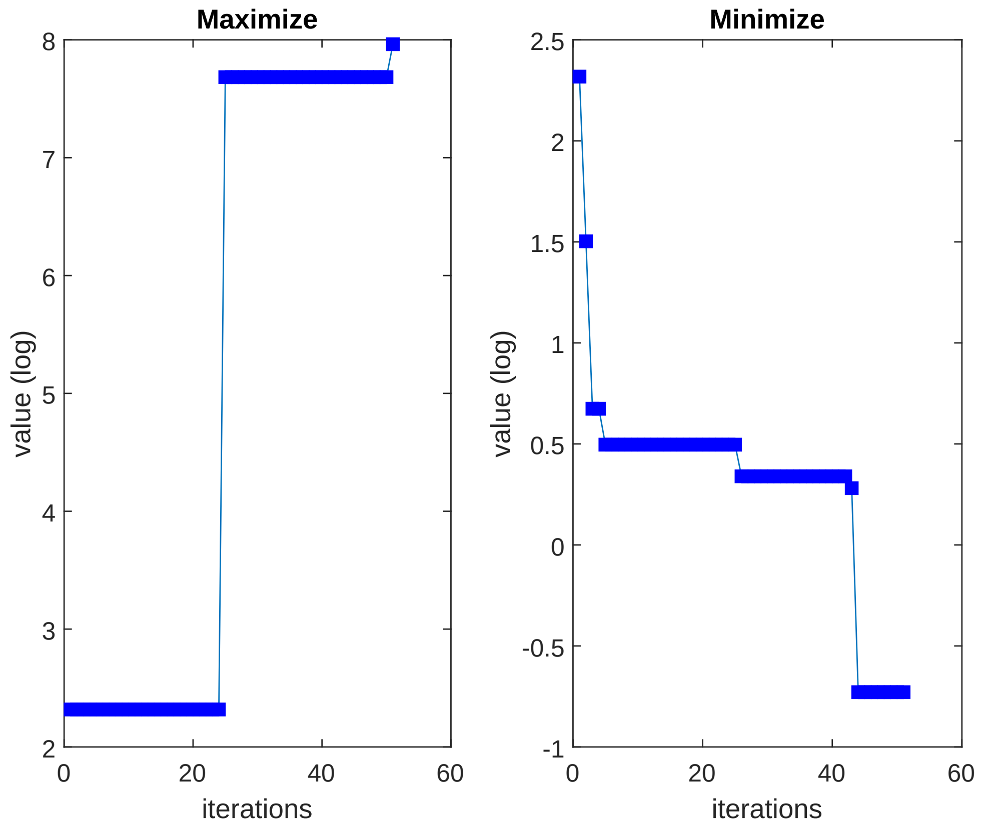

Supplementary Figure 2: The waterfall plot for the parameter k_poolIn. The parameter value of k_poolIn was both minimized and maximized to get the bounds, while still requiring the overall agreement to data to be sufficiently good (determined with a $\chi^2$-test). Parameter k_poolIn is identifiable.

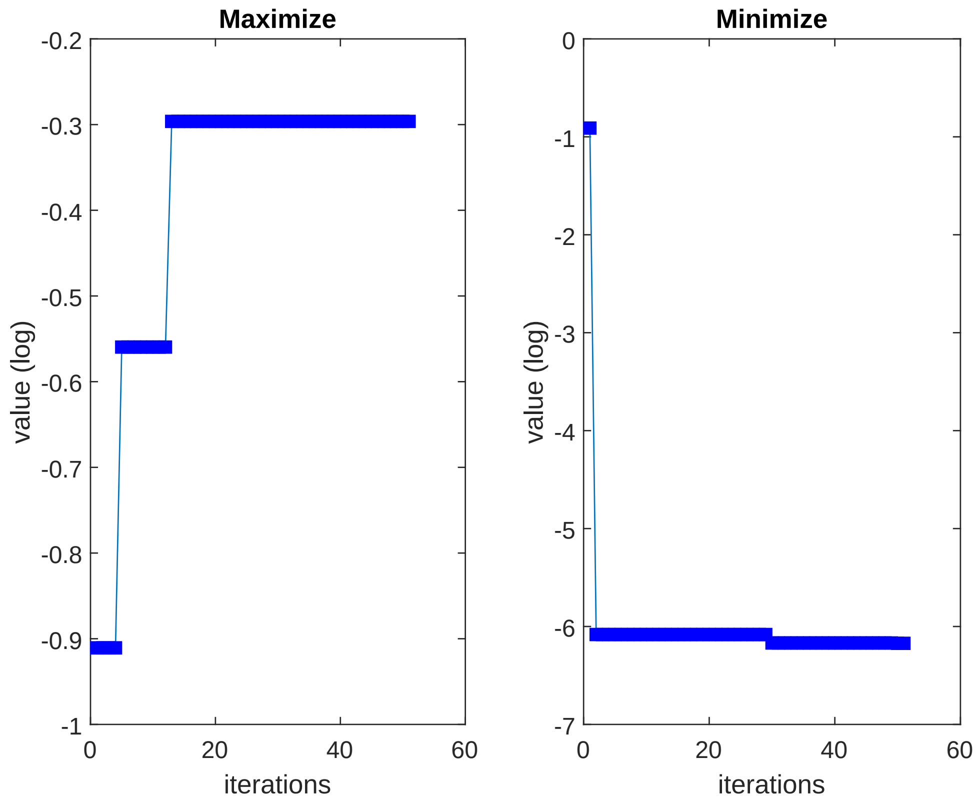

Supplementary Figure 3: The waterfall plot for the parameter k_poolOut. The parameter value of k_poolOut was both minimized and maximized to get the bounds, while still requiring the overall agreement to data to be sufficiently good (determined with a $\chi^2$-test). Parameter k_poolOut is identifiable.

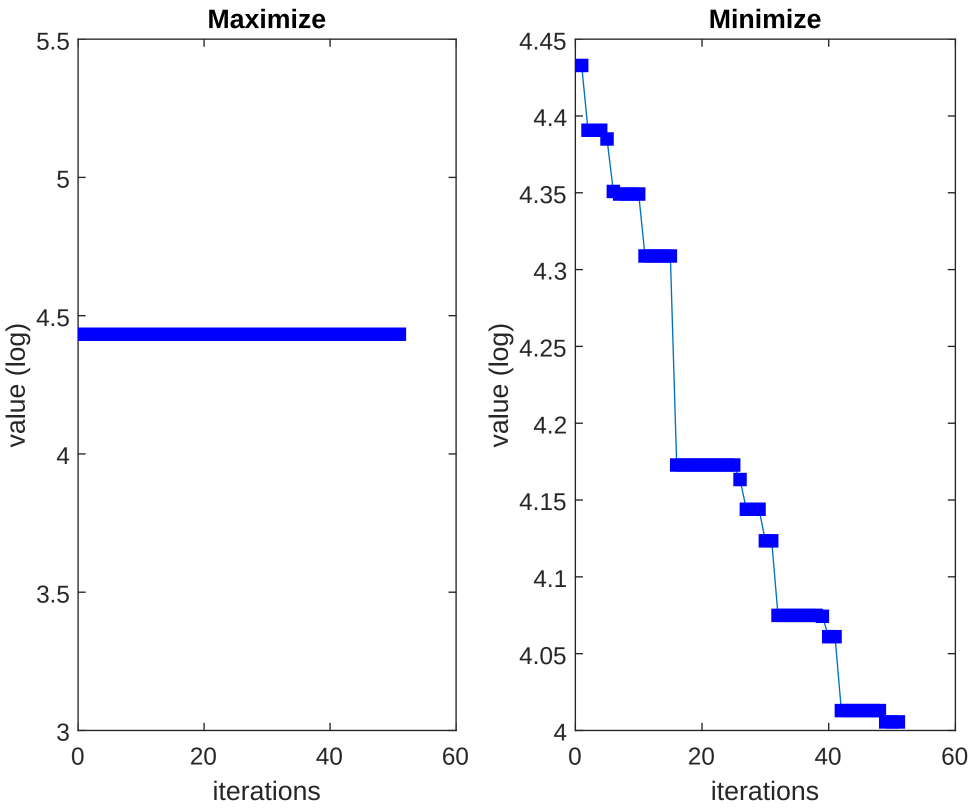

Supplementary Figure 4: The waterfall plot for the parameter k3. The parameter value of k3 was both minimized and maximized to get the bounds, while still requiring the overall agreement to data to be sufficiently good (determined with a $\chi^2$-test). Parameter k3 is identifiable.

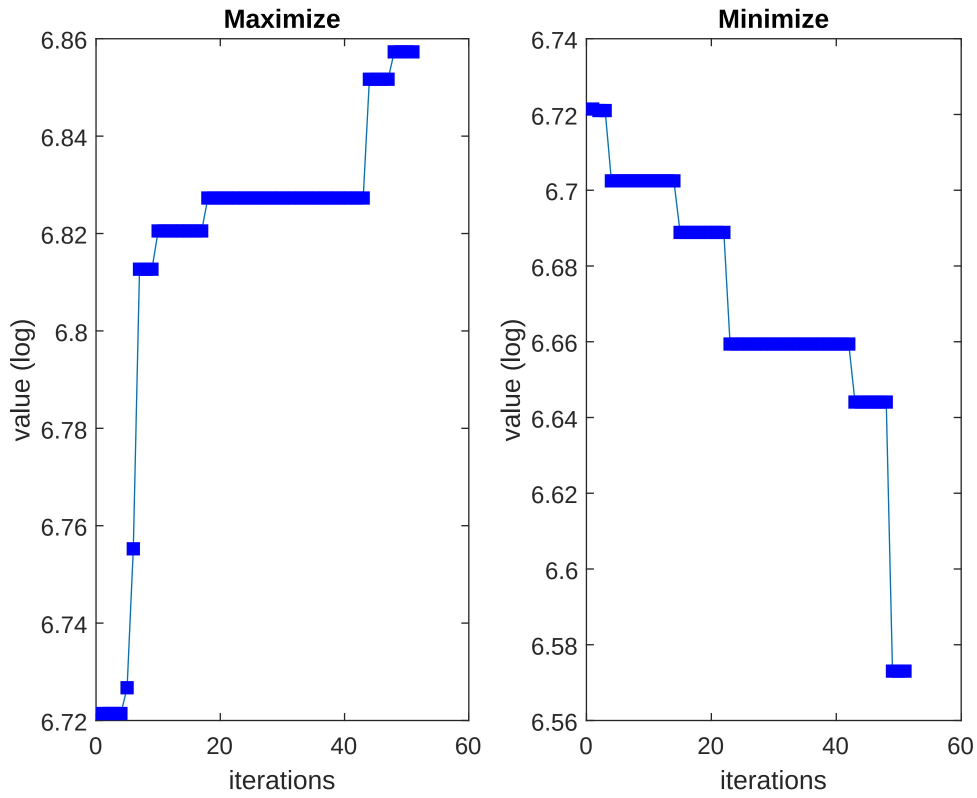

Supplementary Figure 5: The waterfall plot for the parameter k4. The parameter value of k4 was both minimized and maximized to get the bounds, while still requiring the overall agreement to data to be sufficiently good (determined with a $\chi^2$-test). Parameter k4 is identifiable.

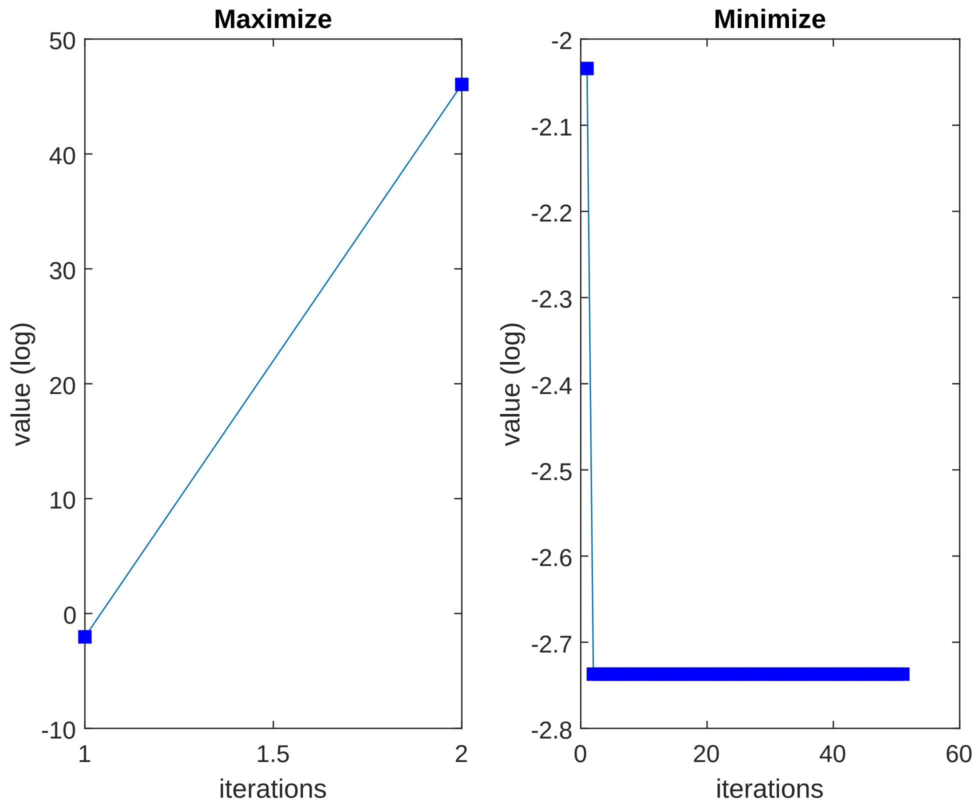

Supplementary Figure 6: The waterfall plot for the parameter k6. The parameter value of k6 was both minimized and maximized to get the bounds, while still requiring the overall agreement to data to be sufficiently good (determined with a $\chi^2$-test). Parameter k6 is (upwards) nonidentifiable.

Notably, parameter k6 is (upwards) nonidentifiable (Supplementary Figure 6). This is because the inclusion of only one data set of acetate elimination in the liver, which has a small impact on the overall objective function value. Due to this small influence, the rate of acetate elimination, governed by k6, can become unlimitedly large. In practice, a very large elimination rate will effectively eliminate all acetate, and thus keep the acetate concentration at 0 over the time series. To limit the elimination rate, additional acetate data would be needed.

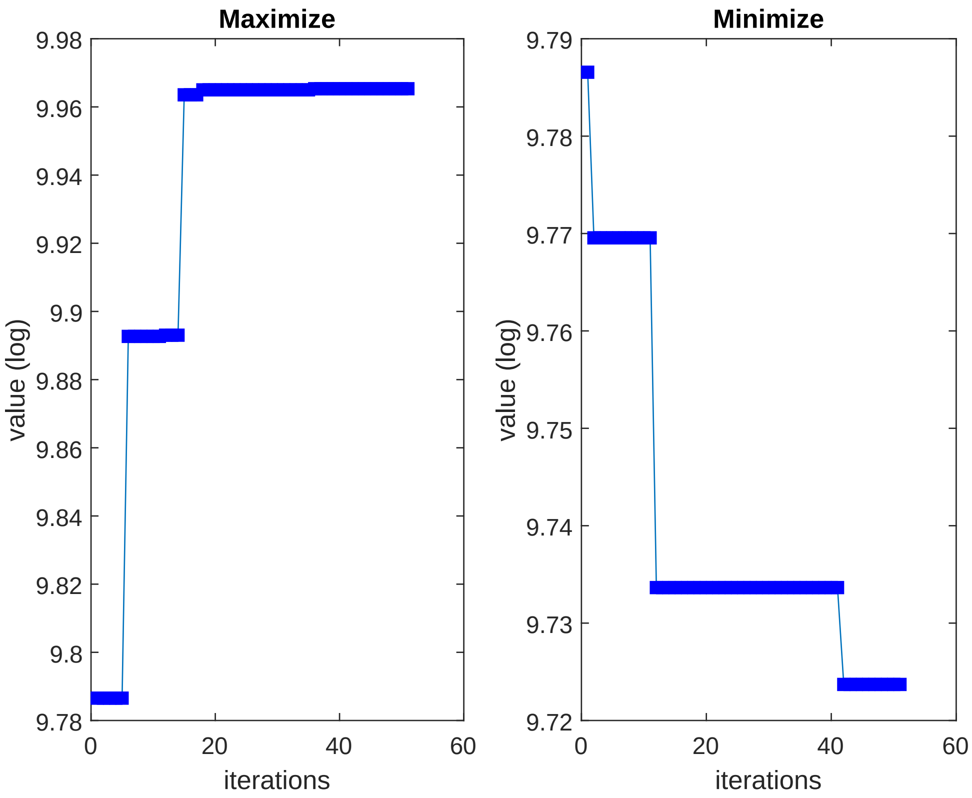

Supplementary Figure 7: The waterfall plot for the parameter km. The parameter value of km was both minimized and maximized to get the bounds, while still requiring the overall agreement to data to be sufficiently good (determined with a $\chi^2$-test). Parameter km is identifiable.

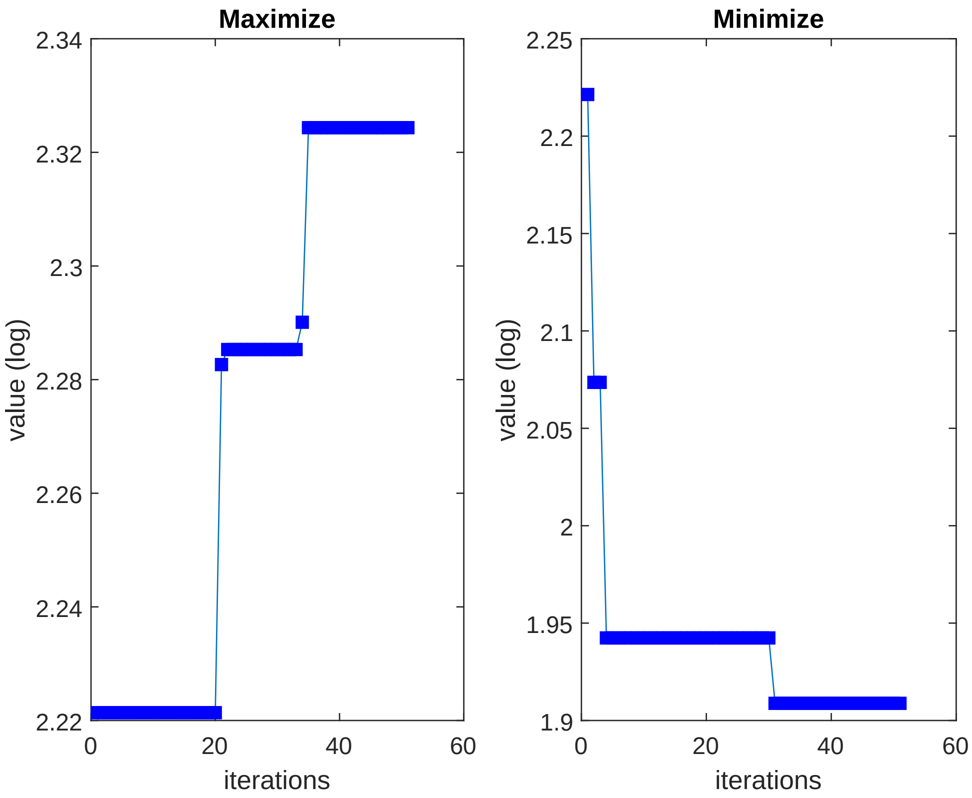

Supplementary Figure 8: The waterfall plot for the parameter kmADH. The parameter value of kmADH was both minimized and maximized to get the bounds, while still requiring the overall agreement to data to be sufficiently good (determined with a $\chi^2$-test). Parameter kmADH is identifiable.

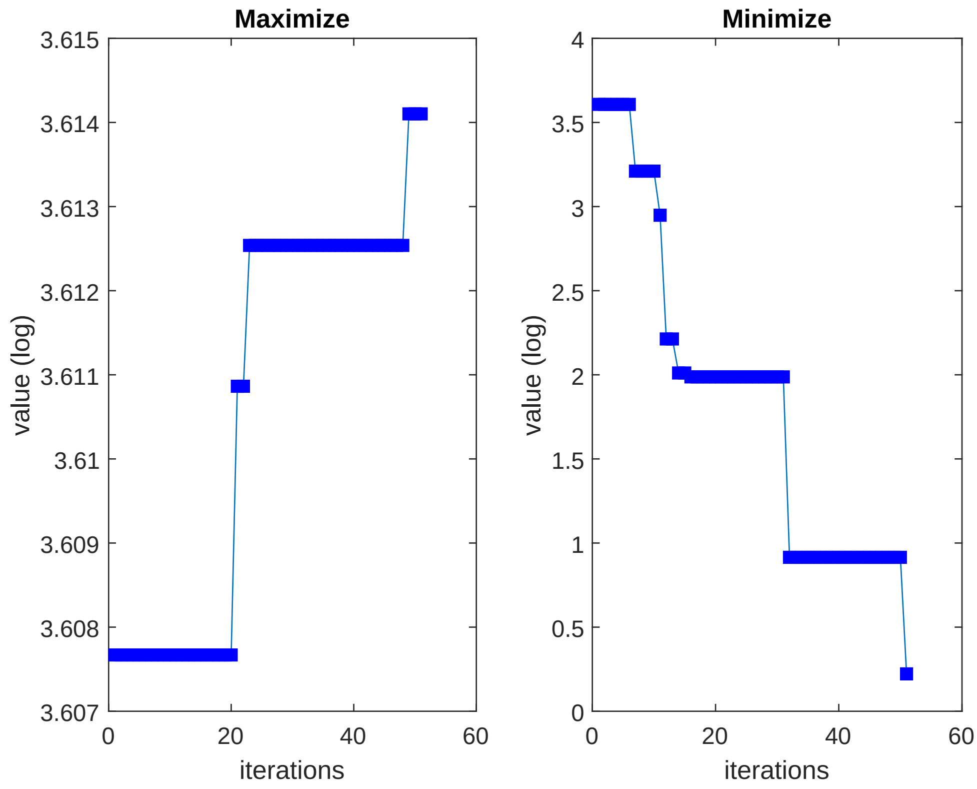

Supplementary Figure 9: The waterfall plot for the parameter kmCYP2E1. The parameter value of kmCYP2E1 was both minimized and maximized to get the bounds, while still requiring the overall agreement to data to be sufficiently good (determined with a $\chi^2$-test). Parameter kmCYP2E1 is identifiable.

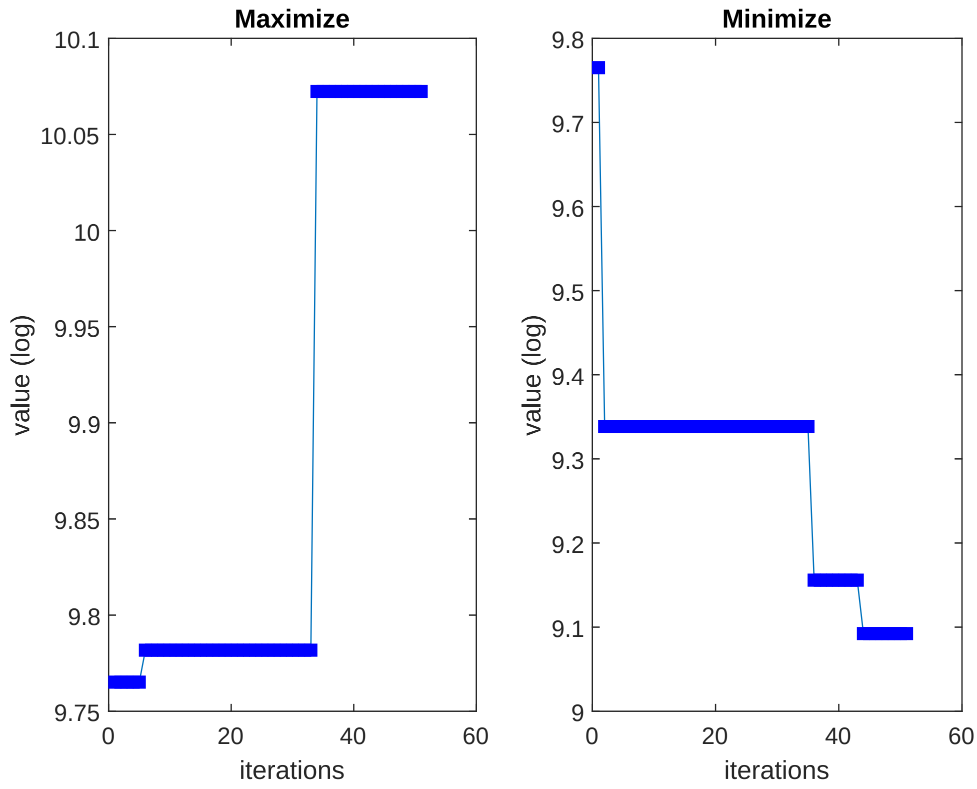

Supplementary Figure 10: The waterfall plot for the parameter kPEth_bind. The parameter value of kPEth_bind was both minimized and maximized to get the bounds, while still requiring the overall agreement to data to be sufficiently good (determined with a $\chi^2$-test). Parameter kPEth_bind is identifiable.

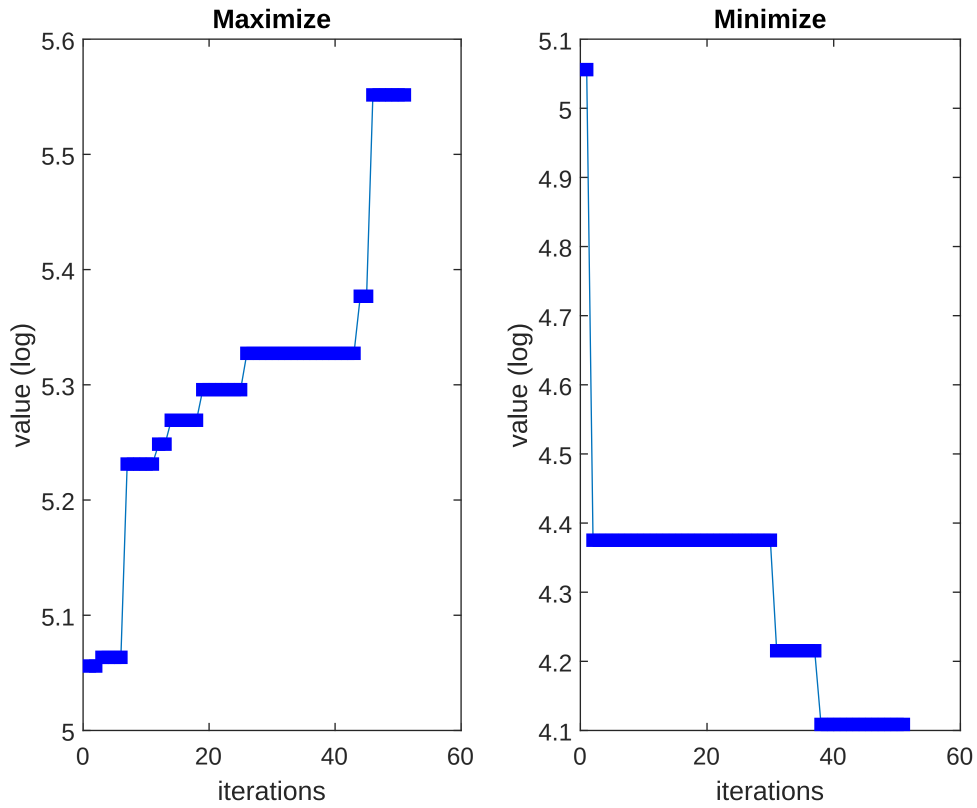

Supplementary Figure 11: The waterfall plot for the parameter kPEth_out. The parameter value of kPEth_out was both minimized and maximized to get the bounds, while still requiring the overall agreement to data to be sufficiently good (determined with a $\chi^2$-test). Parameter kPEth_out is identifiable.

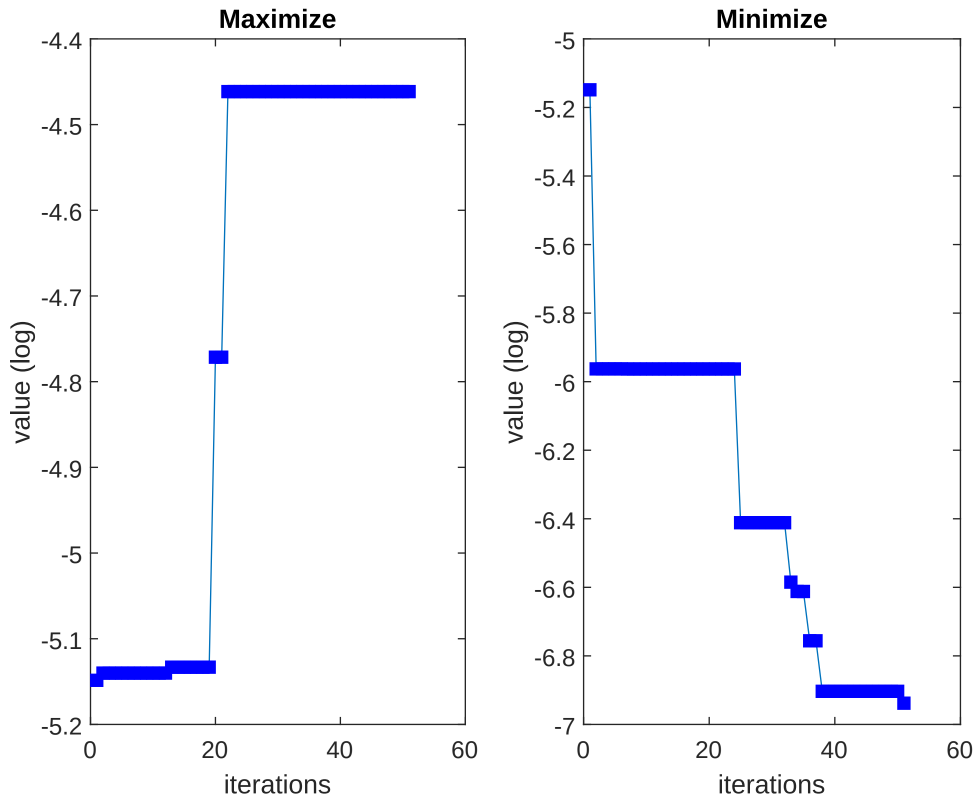

Supplementary Figure 12: The waterfall plot for the parameter kPEth_release. The parameter value of kPEth_release was both minimized and maximized to get the bounds, while still requiring the overall agreement to data to be sufficiently good (determined with a $\chi^2$-test). Parameter kPEth_release is identifiable.

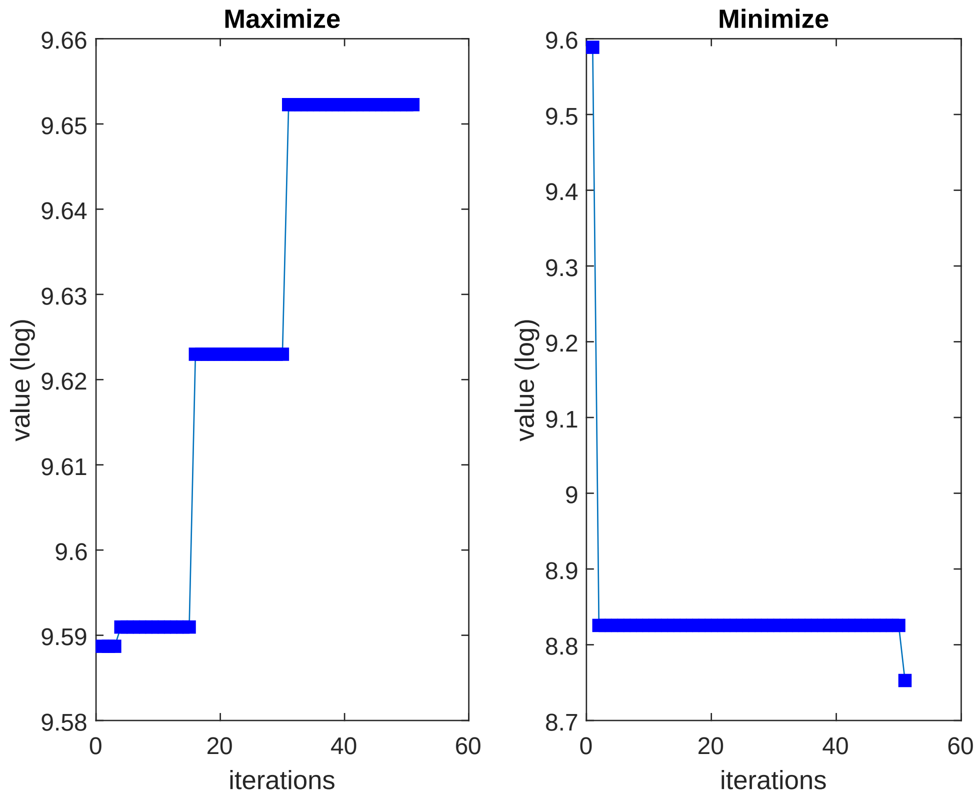

Supplementary Figure 13: The waterfall plot for the parameter kPEth. The parameter value of kPEth was both minimized and maximized to get the bounds, while still requiring the overall agreement to data to be sufficiently good (determined with a $\chi^2$-test). Parameter kPEth is identifiable.

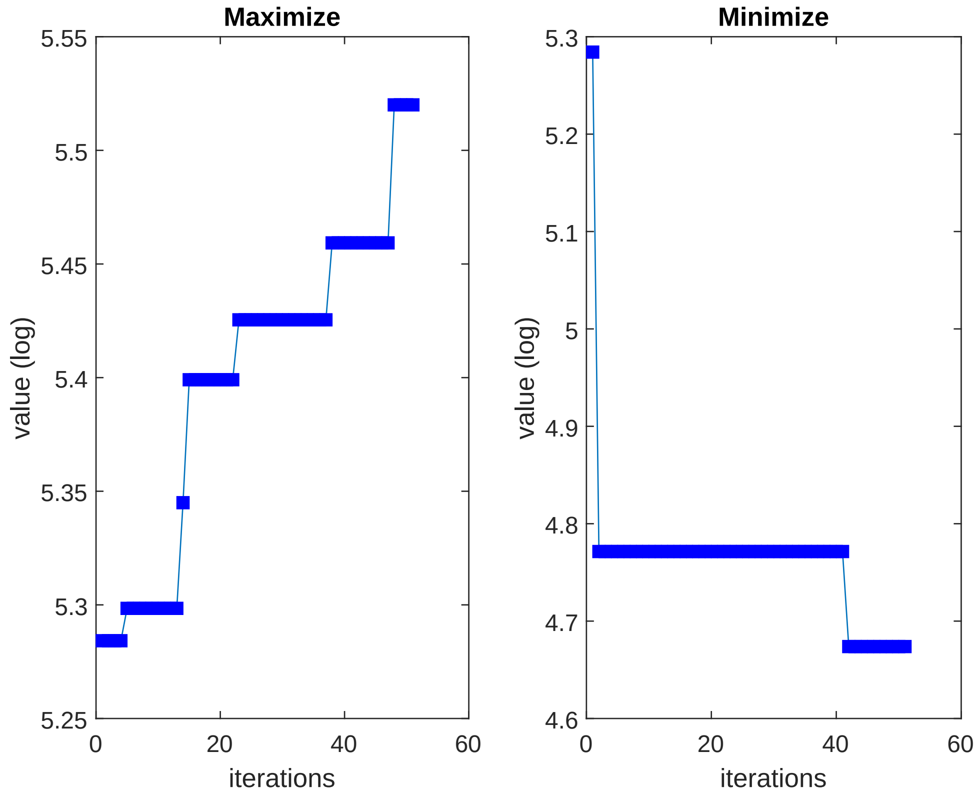

Supplementary Figure 14: The waterfall plot for the parameter PEth_h. The parameter value of PEth_h was both minimized and maximized to get the bounds, while still requiring the overall agreement to data to be sufficiently good (determined with a $\chi^2$-test). Parameter PEth_h is identifiable.

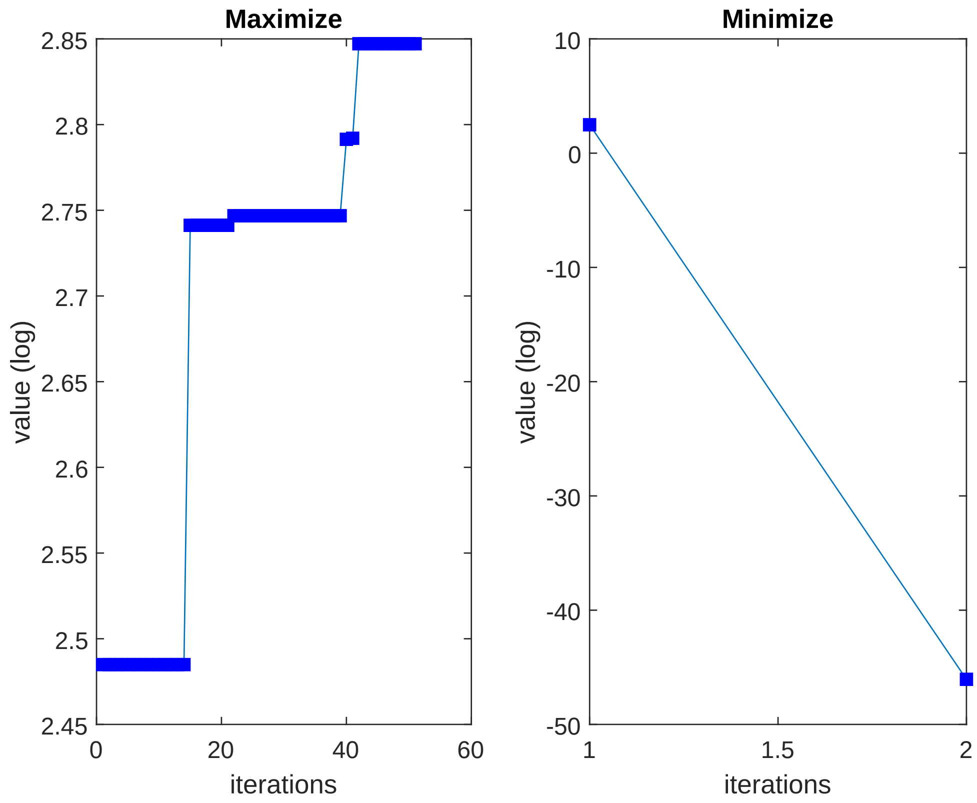

Supplementary Figure 15: The waterfall plot for the parameter PEth_L. The parameter value of PEth_L was both minimized and maximized to get the bounds, while still requiring the overall agreement to data to be sufficiently good (determined with a $\chi^2$-test). Parameter PEth_L is (downwards) nonidentifiable.

PEth_L corresponds to the initial value of PEth in the "low dose" group from the study by Javors et al. [1] and is (downwards) nonidentifiable (Supplementary Figure 15). While it is unlikely that group would have no traces of PEth when starting the study, it is still possible to explain the PEth data which included the group, with a basal level of PEth at 0. Since no basal levels of the low group was given in the study by Javors et al., we cannot know if the basal level was 0 or not.

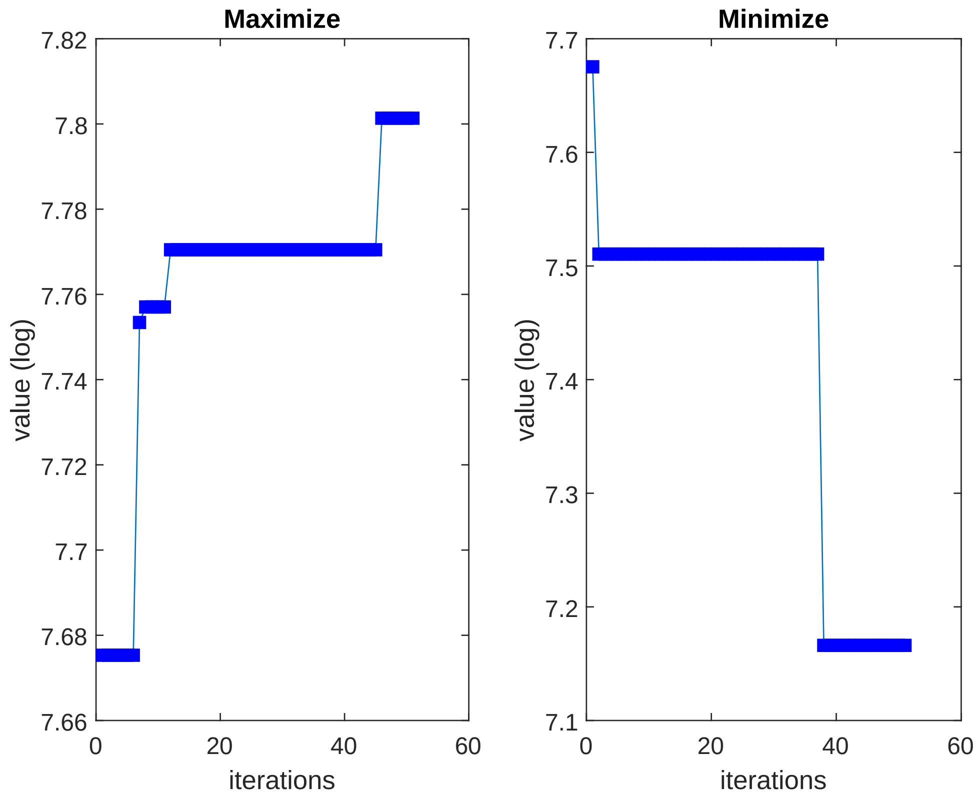

Supplementary Figure 16: The waterfall plot for the parameter Vmax. The parameter value of Vmax was both minimized and maximized to get the bounds, while still requiring the overall agreement to data to be sufficiently good (determined with a $\chi^2$-test). Parameter Vmax is identifiable.

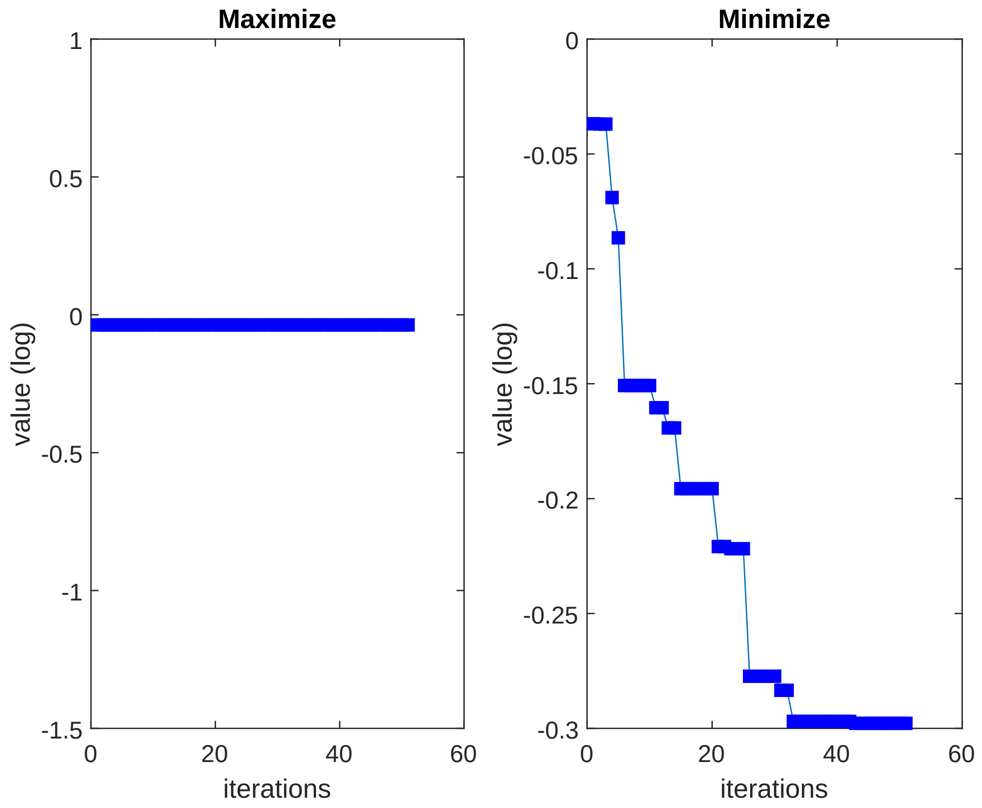

Supplementary Figure 17: The waterfall plot for the parameter VmaxADH. The parameter value of VmaxADH was both minimized and maximized to get the bounds, while still requiring the overall agreement to data to be sufficiently good (determined with a $\chi^2$-test). Parameter VmaxADH is identifiable.

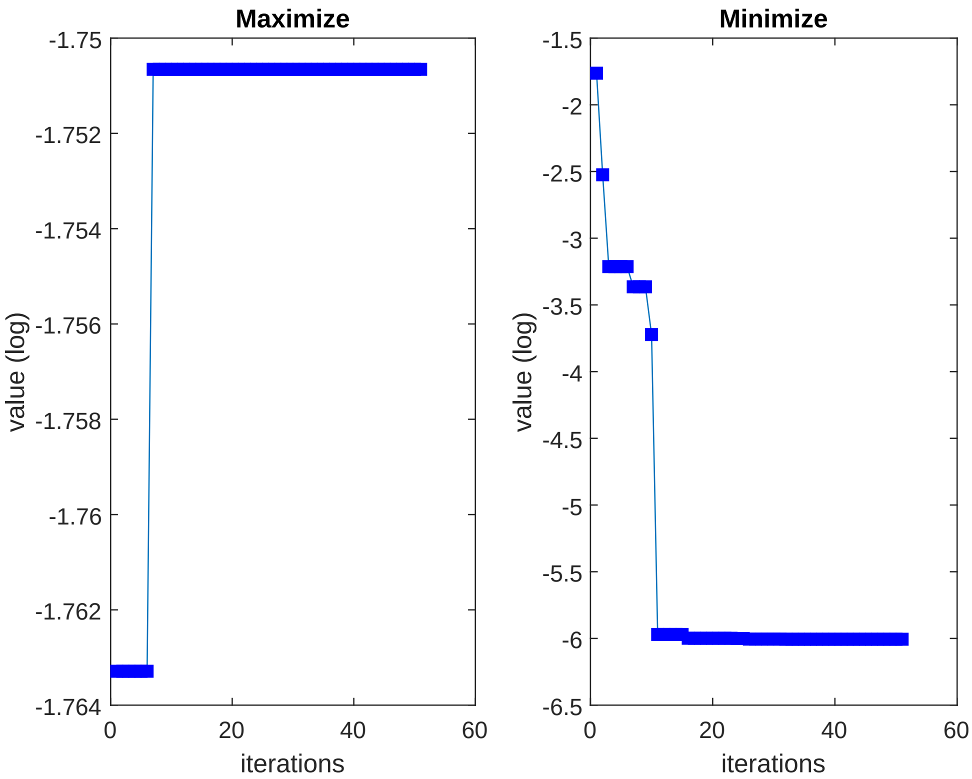

Supplementary Figure 18: The waterfall plot for the parameter VmaxCYP2E1. The parameter value of VmaxCYP2E1 was both minimized and maximized to get the bounds, while still requiring the overall agreement to data to be sufficiently good (determined with a $\chi^2$-test). Parameter VmaxCYP2E1 is identifiable.

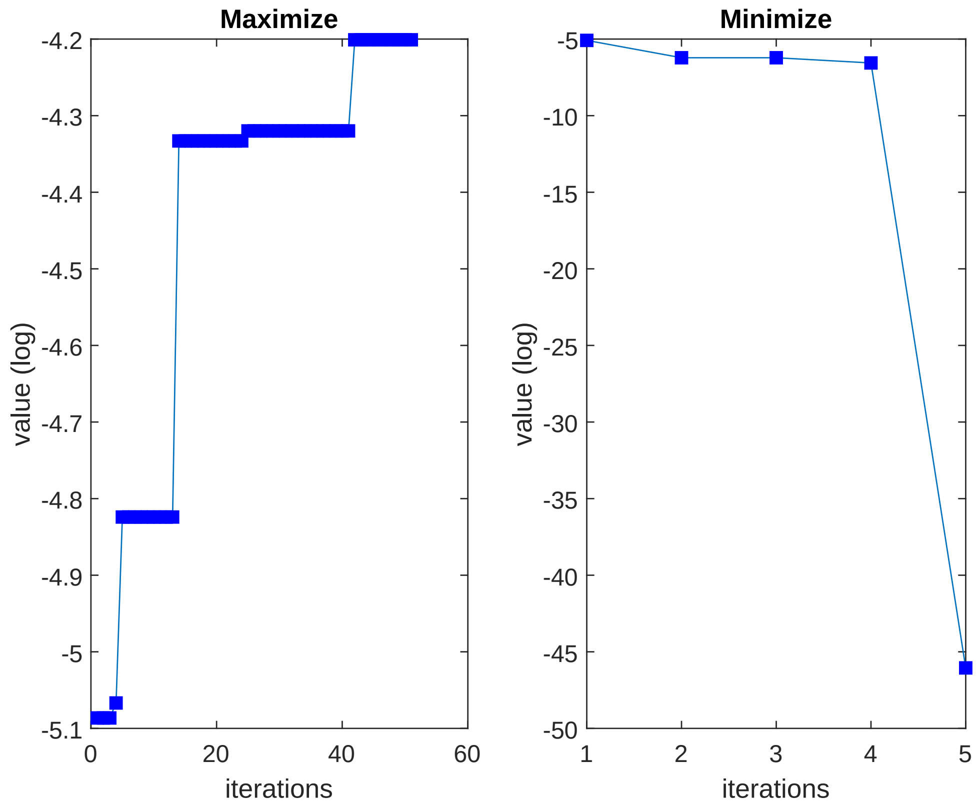

Supplementary Figure 19: The waterfall plot for the parameter k_Kcal_clearance. The parameter value of k_Kcal_clearance was both minimized and maximized to get the bounds, while still requiring the overall agreement to data to be sufficiently good (determined with a $\chi^2$-test). Parameter k_Kcal_clearance is nonidentifiable.

Lastly, k_Kcal_Clearance, which corresponds to the rate of kcal clearance in the stomach, is (downwards) nonidentifiable. However, with k_Kcal_Clearance set to 0, there would not be any clearance of the kcal in the stomach, which is not reasonable, since we know that fasting leads to emptying of the stomach. The reason for why k_Kcal_Clearance can be set to 0 and still provide a sufficiently good agreement between the simulation and the data, is that we in the training only included data on single drinks. However, without the inclusion of a kcal clearance, the kcal would accumulate over time and reduce the response to new drinks, until no response at all would be achieved (Supplementary Figure 20). The inclusion of the parameter leads to more realistic drinking behaviour when consuming multiple drinks (Figure 7 in the main article) and thus better predictions.

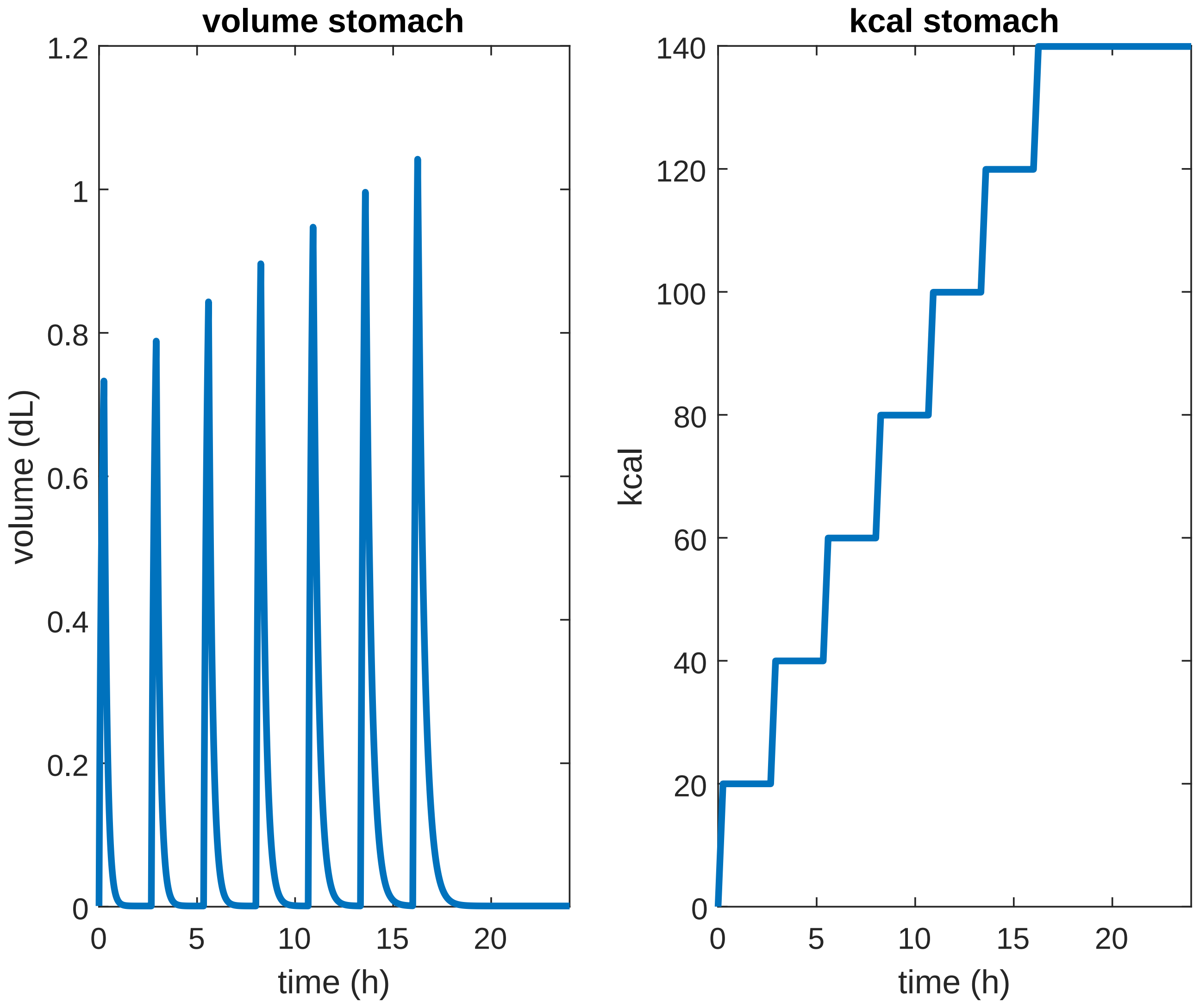

Supplementary Figure 20: Dynamics of consuming multiple drinks without kcal clearance. Here, 7 drinks of wine (15cl) is consumed and the stagnating dynamic is illustrated by A) the volume of liquid in the stomach is increasing even if the drink size is the same, and B) the amount of kcal in the stomach is accumulating with every drink.

### 2 Analysis of the impact of the model inputs on ethanol dynamics

We performed a sensitivity analysis to investigate the impact of the different model inputs on the ethanol dynamics. The sensitivity analysis was performed by varying each input individually, while keeping all other inputs constant. We varied the inputs in a range between a 50% decrease to a 50% increase. As can be seen, inputs affecting the blood volume (height, weight, and sex) have a large impact on the ethanol dynamics (Supplementary Figure 21A-D).

We also varied the activity of the enzymes ADH (Supplementary Figure 21E) and CYP2E1 (Supplementary Figure 21F). Varying the activity of ADH had a larger impact than varying the CYP2E1 activity.

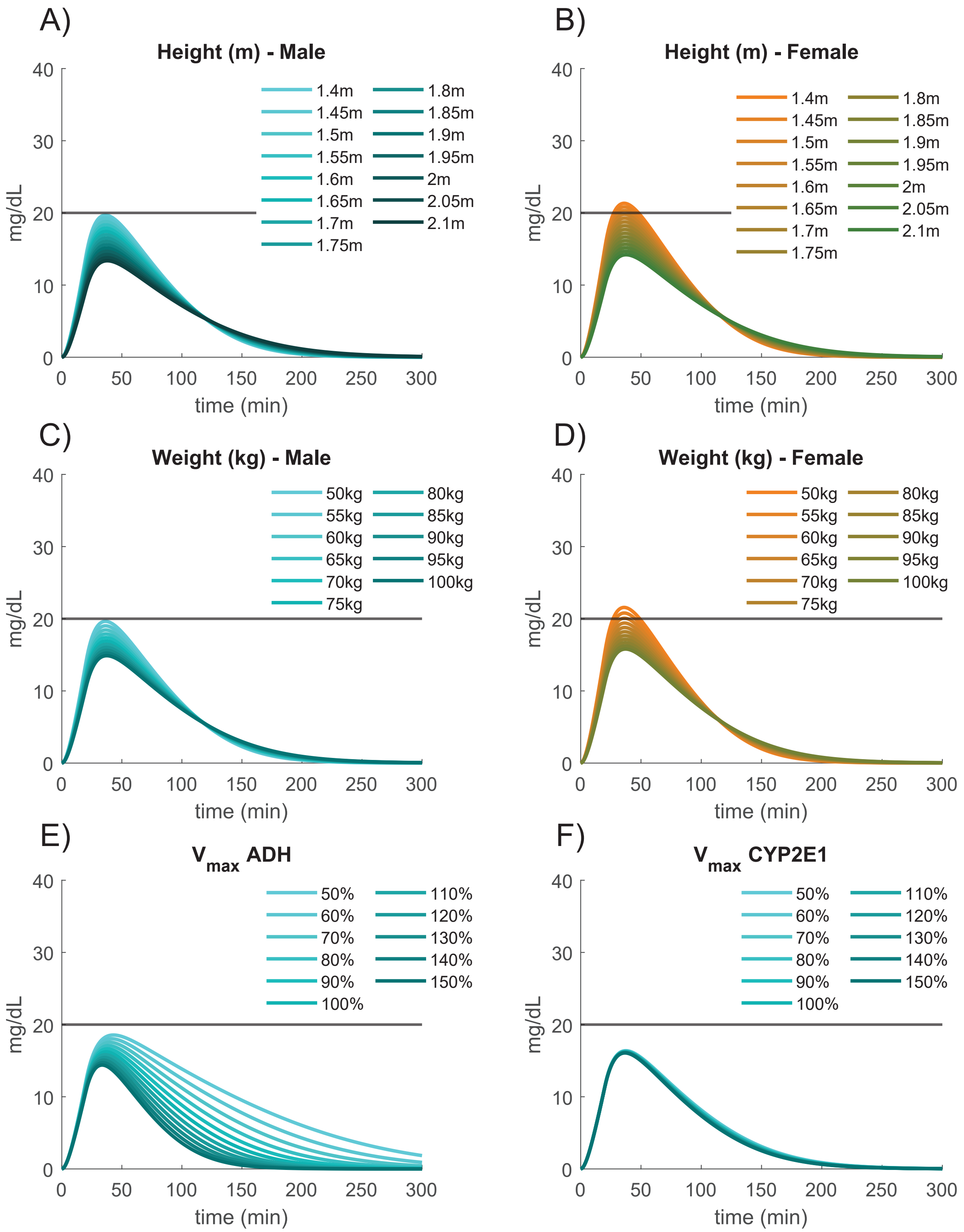

Supplementary Figure 21: The impact of varying different inputs. A) varying the height parameter for men. B) varying the height parameter for women. C) varying the weight parameter for men. D) varying the height parameter for women. E) varying the activity of ADH. F) varying the activity of CYP2E1.

We also tested mixing beverages in different orders (Supplementary Figure 22A) and with different delays (Supplementary Figure 22B). In Supplementary Figure 22A, we can observe that consuming a glass of juice in between two beers reduces the amplitude of the BAC profile compared to consuming the same two beers without a glass of juice. The overall time for the BAC to reach zero is, however, mostly unchanged. The dashed lines illustrate how the BAC profile would look if the second beer was not consumed. From this one can observe that the effect of the juice is the greatest if consumed before the alcoholic beverage. This stems from the dynamics of the gastric emptying. As the addition of juice increases the caloric content the rate of gastric emptying decreases. However, if one drinks the beer before the juice, most of the beer has already emptied before the juice is added. This is the reason the BAC clearance is just slightly faster for Beer + Juice (dashed line).

Moreover, Supplementary Figure 22B showcases how the timing of the juice affects the BAC profile. Similar to Supplementary Figure 22A, if one consumes juice after a shot the amplitude is reduced and the overall clearance time is mostly unaffected. Consuming the juice before the shot gives a lower peak and a lower slope. This effect is greater the closer the juice is consumed to the shot. If the juice is consumed after the shot, the slope remains unaffected, and the peak amplitude is lowered. This shows that by integrating the dynamics of gastric emptying our model can replicate real-life behaviors seen from consuming different beverages. These predictions are in line with previous experiments, Mitchell et al. [2], showing that the same alcohol content consumed in different beverages gives an amplitude reduction but mostly unaffected clearance time. Our models suggest that similar effects can be achieved by mixing the alcohol content in the stomach, arguing for the effect of mixing in non-alcoholic beverages to reduce the peak BAC amplitude and slowing down drunkenness.

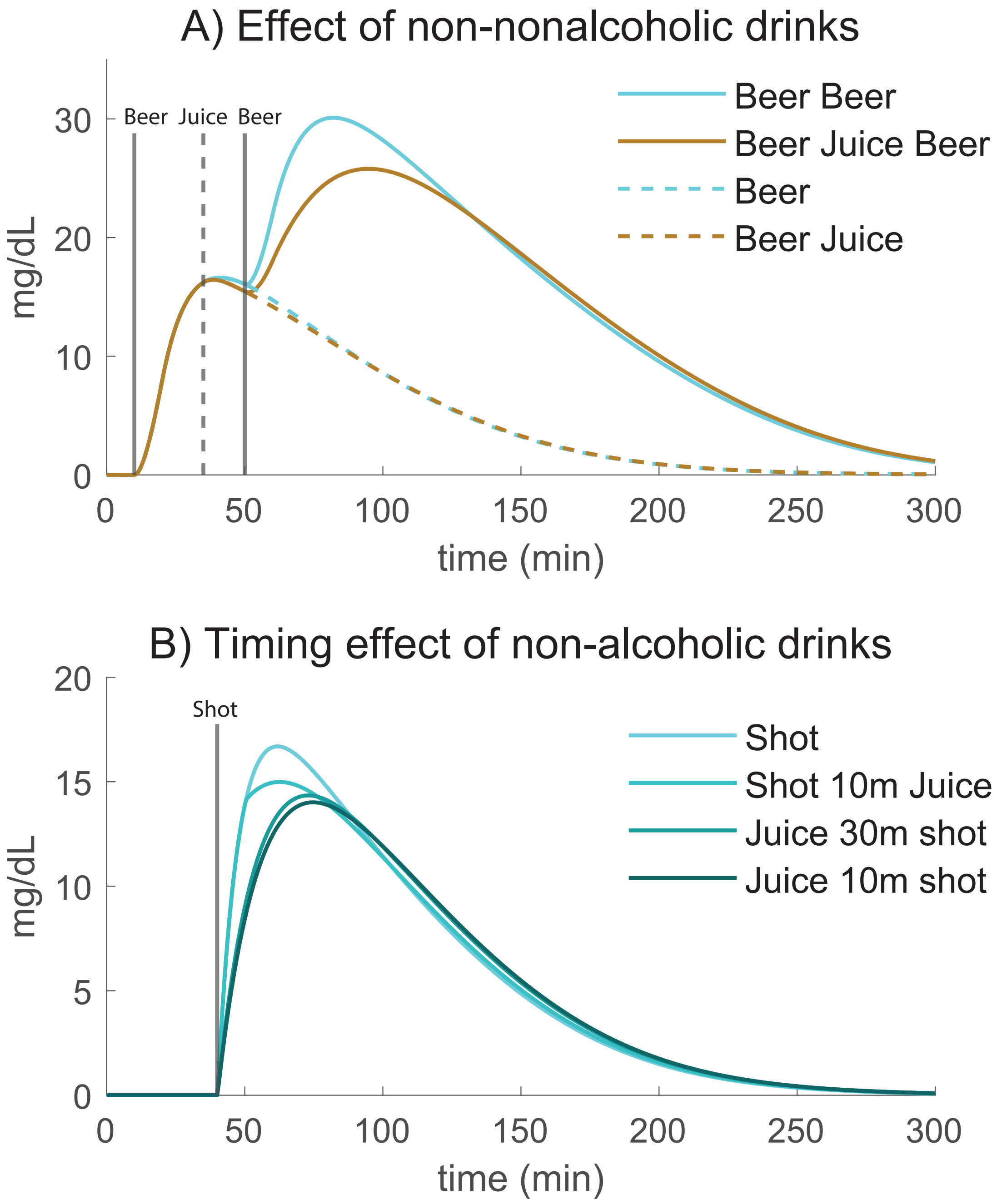

Supplementary Figure 22: The impact of varying the timing and order of drinks. A) The effect of consuming a glass of juice (0.33L, 40kcal/dL) between two beers (0.33L, 5.1v/v%, 13kcal/dL) on the BAC profile. The beers are consumed over 10 min, starting at the solid x-lines (t=10 & t=50), and the juice is consumed over 10 min, starting at the dashed x-line (t=35). The dashed lines show how the BAC profile would have been if the second beer was excluded. B) The effect of changing the timing of consuming the non-alcoholic drink. Here, a shot (0.04L,40v/v%) was consumed over 15 sec, starting at the x-line (t=40). One glass of juice (0.33L, 40kcal/dL) was consumed over 10 min with different timing to the shot, i) no juice, ii) juice 10 min after the shot, iii) juice 30 min before the shot, and iv) juice 10 min before the shot.

### 3 Input estimations

This section presents all the inferred values that has been estimated for the different data sets and how this was performed.

#### 3.1 Methods for estimating values

##### 3.1.1 Estimating the height of the participants when no height where given

We used databases to estimate the height, using the national averages in from the country of origin of the original paper, and if possible also the national average for the year the paper was published. All such values are given in the tables below.

##### 3.1.2 Calculating weight from BMI

If the BMI of the participants were given in the original paper but not the weight, then we calculated the weight based on the height of the participants and the BMI, using the definition of BMI.

$$weight = BMI \cdot height^2 \tag{1}$$

##### 3.1.3 Estimating drink volume from g ethanol per kg body weight and concentration

In many studies, the full details of the drinks were not given in the paper. The amount of alcohol given was often defined as g ethanol per kg body weight of the participants. Thus we could calculate the total grams of alcohol given by multiplying the ethanol g/kg body weight (X) with the body weight:

$$mass_{ethanol} = X \cdot weight \tag{2}$$

Using the density of alcohol (g/ml), we could calculate the total volume (ml) of ethanol in the drink:

$$V_{ethanol} = mass_{ethanol}/0.7891 \tag{3}$$

Knowing the final volume of ethanol in the drink, we could calculate the total volume using the dilution equation:

$$c_{ethanol} \cdot V_{ethanol} = c_{drink} \cdot V_{drink} \tag{4}$$

Here we assume that the concentration of ethanol is 100%, thus $c_{ethanol} = 1$ and:

$$V_{drink} = vol_{ethanol}/c_{drink} \tag{5}$$

If no concentration of the final drink was given in the original paper, then we assumed the concentration to be 10% (v/v%).

##### 3.1.4 Average sex in a group

In some datasets the participants were both men and women. From the perspective of the model, the major difference between individual were driven by the differences in total blood volume. Thus, if the dataset contained both men and women, we needed to construct an average person (either man or woman) but with the average blood volume. For this, we first determined if men or women were the most common sex in the group, and then we calculated the blood volumes for the subsection of either sex, and then constructed an average individual. We calculated the blood volume using the Nadler equation [3]. For men blood volume was estimated using, where the height is given in meters:

$$vol_{blood,male} = (0.3669 \cdot height_{male}^3 + 0.03219 \cdot weight_{male} + 0.6041) \tag{6}$$

And for women:

$$vol_{blood,female} = (0.3561 \cdot height_{female}^3 + 0.03308 \cdot weight_{female} + 0.1833) \tag{7}$$

We then calculated the ratio of women to men according to, where sex=1 if man, and sex=0 if woman:

$$sex = n_{males}/(n_{females} + n_{males}) \tag{8}$$

We then calculated the average blood volume within the group:

$$vol_{blood} = (sex) \cdot vol_{blood,male} + (1 - sex) \cdot vol_{blood,female} \tag{9}$$

From the average blood volume, we reverse calculated an individual with the same blood volume. If there were more men in the group we assumed the height to be the average height within the men in the group, and then calculated the weight using the Nadler equation according to:

$$weight = (vol_{blood} - 0.6041 - 0.3669 \cdot height_{male}^3)/0.03219 \tag{10}$$

If there were more women than men, we instead used the average height of the women, and calculated the weight according to:

$$weight = (vol_{blood} - 0.1833 - 0.3561 \cdot height_{female}^3)/0.03308 \tag{11}$$

##### 3.1.5 Estimating calories within the drink

If the authors of the original papers had not disclosed the caloric content of the drinks, we approximated the calories based on the drink type. Note that the calories of the drink excludes the calories from the ethanol.

#### 3.2 Information for each dataset

Below are the values used for the different experiments. If the value were given in the paper, the value in the “Given values” column was used, and if not, we inferred the value as described above and used the value in the “Inferred values” column. The different inputs correspond to:

- sex: the most frequent sex of the participants. 0 corresponds to women, and 1 corresponds to men.

- height: the height in meters
- weight: the weight in kg
- BMI: the body mass index. Was used to infer the weight if no information on the weight was given.
- volume: The volume of the drink consumed
- concentration: The concentration of the drink consumed (in v/v%)
- drink kcal: the amount of kilocalories in the drink, excluding the calories from ethanol
- drink length: under how long time (minutes) the drink was consumed
- food kcal: the amount of kilocalories in food consumed
- time of meal: the time when the meal was consumed (relative to the start of the drink)

Below we give the values for each dataset/experiment.

##### 3.2.1 Okabe 500 ml Water

| **Input** | **Given values** | **Inferred values** |
|---|---|---|
| Sex | 1 | – |
| Height | 1.71 | – |
| Weight | 65 | – |
| BMI | 22 | – |
| Volume | 0.5 | – |
| Concentration | 0.0 | – |
| Drink kcal | 0 | – |
| Drink length | 3 | – |
| Food kcal | 0 | – |
| Time of meal | 0 | – |

Supplementary Table 1: **Input values for Okabe 500 ml water**

###### 3.2.2 Okabe 500 ml orange juice

| Input | Given values | Inferred values |
|---|---|---|
| Sex | 1 | – |
| Height | 1.71 | – |
| Weight | 65 | – |
| BMI | 22 | – |
| Volume | 0.5 | – |
| Concentration | 0.0 | – |
| Drink kcal | 220 | – |
| Drink length | 3 | – |
| Food kcal | 0 | – |
| Time of meal | 0 | – |

Supplementary Table 2: **Input values for Okabe 500 ml orange juice**

###### 3.2.3 Okabe 500 ml milk water

| Input | Given values | Inferred values |
|---|---|---|
| Sex | 1 | – |
| Height | 1.71 | – |
| Weight | 65 | – |
| BMI | 22 | – |
| Volume | 0.5 | – |
| Concentration | 0.0 | – |
| Drink kcal | 220 | – |
| Drink length | 3 | – |
| Food kcal | 0 | – |
| Time of meal | 0 | – |

Supplementary Table 3: **Input values for Okabe 500 ml milk water**

##### 3.2.4 Okabe 500 ml orange juice and syrup

| Input | Given values | Inferred values |
|---|---|---|
| Sex | 1 | – |
| Height | 1.71 | – |
| Weight | 65 | – |
| BMI | 22 | – |
| Volume | 0.5 | – |
| Concentration | 0.0 | – |
| Drink kcal | 330 | – |
| Drink length | 3 | – |
| Food kcal | 0 | – |
| Time of meal | 0 | – |

Supplementary Table 4: **Input values for Okabe 500 ml orange juice and syrup**

##### 3.2.5 Okabe 500 ml milk

| Input | Given values | Inferred values |
|---|---|---|
| Sex | 1 | – |
| Height | 1.71 | – |
| Weight | 65 | – |
| BMI | 22 | – |
| Volume | 0.5 | – |
| Concentration | 0.0 | – |
| Drink kcal | 330 | – |
| Drink length | 3 | – |
| Food kcal | 0 | – |
| Time of meal | 0 | – |

Supplementary Table 5: **Input values for Okabe 500 ml milk**

###### 3.2.6 Okabe 150 ml water

| Input | Given values | Inferred values |
|---|---|---|
| Sex | 1 | – |
| Height | 1.712 | – |
| Weight | 68.4 | – |
| BMI | 23 | – |
| Volume | 0.15 | – |
| Concentration | 0.0 | – |
| Drink kcal | 0 | – |
| Drink length | 3 | – |
| Food kcal | 0 | – |
| Time of meal | 0 | – |

Supplementary Table 6: **Input values for Okabe 150 ml water**

###### 3.2.7 Okabe 150 ml glucose

| Input | Given values | Inferred values |
|---|---|---|
| Sex | 1 | – |
| Height | 1.712 | – |
| Weight | 68.4 | – |
| BMI | 23 | – |
| Volume | 0.15 | – |
| Concentration | 0.0 | – |
| Drink kcal | 67 | – |
| Drink length | 3 | – |
| Food kcal | 0 | – |
| Time of meal | 0 | – |

Supplementary Table 7: **Input values for Okabe 150 ml glucose**

###### 3.2.8 Okabe 150 ml whiskey

| Input | Given values | Inferred values |
|---|---|---|
| Sex | 1 | – |
| Height | 1.712 | – |
| Weight | 68.4 | – |
| BMI | 23 | – |
| Volume | 0.15 | – |
| Concentration | 0.0 | – |
| Drink kcal | 0 | – |
| Drink length | 3 | – |
| Food kcal | 0 | – |
| Time of meal | 0 | – |

Supplementary Table 8: **Input values for Okabe 150 ml whiskey**

###### 3.2.9 Okabe 200 ml uniform glucose

| Input | Given values | Inferred values |
|---|---|---|
| Sex | 1 | – |
| Height | 1.69 | – |
| Weight | 60.0 | – |
| BMI | 21 | – |
| Volume | 0.2 | – |
| Concentration | 0.0 | – |
| Drink kcal | 200 | – |
| Drink length | 3 | – |
| Food kcal | 0 | – |
| Time of meal | 0 | – |

Supplementary Table 9: **Input values for Okabe 200 ml uniform glucose**

###### 3.2.10 Okabe 400 ml uniform glucose

| Input | Given values | Inferred values |
|---|---|---|
| Sex | 1 | – |
| Height | 1.69 | – |
| Weight | 60.0 | – |
| BMI | 21 | – |
| Volume | 0.4 | – |
| Concentration | 0.0 | – |
| Drink kcal | 200 | – |
| Drink length | 3 | – |
| Food kcal | 0 | – |
| Time of meal | 0 | – |

Supplementary Table 10: **Input values for Okabe 400 ml uniform glucose**

###### 3.2.11 Okabe 600 ml uniform glucose

| Input | Given values | Inferred values |
|---|---|---|
| Sex | 1 | – |
| Height | 1.69 | – |
| Weight | 60.0 | – |
| BMI | 21 | – |
| Volume | 0.6 | – |
| Concentration | 0.0 | – |
| Drink kcal | 200 | – |
| Drink length | 3 | – |
| Food kcal | 0 | – |
| Time of meal | 0 | – |

Supplementary Table 11: **Input values for Okabe 600 ml uniform glucose**

##### 3.2.12 Mitchell beer

| Input | Given values | Inferred values |
|---|---|---|
| Sex | 1 | – |
| Height | – | 1.77 |
| Weight | 82.66 | – |
| BMI | 26.35 | – |
| Volume | – | 1.027 |
| Concentration | 5.1 | – |
| Drink kcal | – | 133.33 |
| Drink length | 20 | – |
| Food kcal | 0 | – |
| Time of meal | 0 | – |

Supplementary Table 12: **Input values for Mitchell beer**

##### 3.2.13 Mitchell wine

| Input | Given values | Inferred values |
|---|---|---|
| Sex | 1 | – |
| Height | – | 1.77 |
| Weight | 82.66 | – |
| BMI | 26.35 | – |
| Volume | – | 0.419 |
| Concentration | 12.5 | – |
| Drink kcal | – | 55.83 |
| Drink length | 20 | – |
| Food kcal | 0 | – |
| Time of meal | 0 | – |

Supplementary Table 13: **Input values for Mitchell wine**

###### 3.2.14 Mitchell spirit

| Input | Given values | Inferred values |
|---|---|---|
| Sex | 1 | – |
| Height | – | 1.77 |
| Weight | 82.66 | – |
| BMI | 26.35 | – |
| Volume | – | 0.2618 |
| Concentration | 20.0 | – |
| Drink kcal | – | 43.18 |
| Drink length | 20 | – |
| Food kcal | 0 | – |
| Time of meal | 0 | – |

Supplementary Table 14: **Input values for Mitchell spirit**

###### 3.2.15 Jones EtOH and acetaldehyde

| Input | Given values | Inferred values |
|---|---|---|
| Sex | 1 | – |
| Height | – | 1.79 |
| Weight | 74.5 | – |
| BMI | – | 23.25 |
| Volume | 0.25 | – |
| Concentration | – | 9.44 |
| Drink kcal | – | 90.56 |
| Drink length | 5 | – |
| Food kcal | 0 | – |
| Time of meal | 0 | – |

Supplementary Table 15: **Input values for Jones EtOH and acetaldehyde**

##### 3.2.16 Jones fasting

| Input | Given values | Inferred values |
|---|---|---|
| Sex | 1 | – |
| Height | 1.83 | – |
| Weight | 75.8 | – |
| BMI | – | 22.63 |
| Volume | – | 0.1441 |
| Concentration | 20 | 9.44 |
| Drink kcal | – | 51.877 |
| Drink length | 15 | – |
| Food kcal | 0 | – |
| Time of meal | 0 | – |

Supplementary Table 16: **Input values for Jones fasting**

##### 3.2.17 Jones food

| Input | Given values | Inferred values |
|---|---|---|
| Sex | 1 | – |
| Height | 1.83 | – |
| Weight | 75.8 | – |
| BMI | – | 22.63 |
| Volume | – | 0.1441 |
| Concentration | 20 | 9.44 |
| Drink kcal | – | 51.877 |
| Drink length | 15 | – |
| Food kcal | 700 | – |
| Time of meal | -20 | – |

Supplementary Table 17: **Input values for Jones food**

###### 3.2.18 Sarkola

| **Input** | **Given values** | **Inferred values** |
|---|---|---|
| Sex | – | 0 |
| Height | – | 1.66 |
| Weight | – | 75.01 |
| BMI | – | 27.22 |
| Volume | – | 0.475 |
| Concentration | 10 | – |
| Drink kcal | – | 149.61 |
| Drink length | 15 | – |
| Food kcal | 0 | – |
| Time of meal | 0 | – |

Supplementary Table 18: **Input values for Sarkola**

###### 3.2.19 Kechagias fasting

| **Input** | **Given values** | **Inferred values** |
|---|---|---|
| Sex | 1 | – |
| Height | – | 1.80 |
| Weight | 81.9 | – |
| BMI | – | 25.28 |
| Volume | – | 0.156 |
| Concentration | 20 | – |
| Drink kcal | – | 56.188 |
| Drink length | 5 | – |
| Food kcal | 0 | – |
| Time of meal | 0 | – |

Supplementary Table 19: **Input values for Kechagias fasting**

###### 3.2.20 Kechagias breakfast

| Input | Given values | Inferred values |
|---|---|---|
| Sex | 1 | – |
| Height | – | 1.80 |
| Weight | 81.9 | – |
| BMI | – | 25.28 |
| Volume | – | 0.156 |
| Concentration | 20 | – |
| Drink kcal | – | 56.188 |
| Drink length | 5 | – |
| Food kcal | 761.9 | – |
| Time of meal | -60 | – |

Supplementary Table 20: **Input values for Kechagias breakfast**

###### 3.2.21 Javors low dose

| Input | Given values | Inferred values |
|---|---|---|
| Sex | – | 1 |
| Height | – | 1.78 |
| Weight | 58.92 | – |
| BMI | – | 18.60 |
| Volume | 0.70976 | – |
| Concentration | – | 3.245 |
| Drink kcal | – | 242.05 |
| Drink length | 15 | – |
| Food kcal | 0 | – |
| Time of meal | 0 | – |

Supplementary Table 21: **Input values for Javors low dose**

###### 3.2.22 Javors high dose

| Input | Given values | Inferred values |
|---|---|---|
| Sex | – | 1 |
| Height | – | 1.78 |
| Weight | 59.23 | – |
| BMI | – | 18.70 |
| Volume | 0.70976 | – |
| Concentration | – | 6.41 |
| Drink kcal | – | 235.85 |
| Drink length | 15 | – |
| Food kcal | 0 | – |
| Time of meal | 0 | – |

Supplementary Table 22: **Input values for Javors high dose**

###### 3.2.23 Frezza woman

| Input | Given values | Inferred values |
|---|---|---|
| Sex | 0 | – |
| Height | – | 1.61 |
| Weight | – | 59.62 |
| BMI | – | 23 |
| Volume | – | 0.1889 |
| Concentration | – | 12 |
| Drink kcal | – | 33.25 |
| Drink length | 10 | – |
| Food kcal | 555 | – |
| Time of meal | -60 | – |

Supplementary Table 23: **Input values for Frezza woman**

###### 3.2.24 Frezza men

| Input | Given values | Inferred values |
|---|---|---|
| Sex | 1 | – |
| Height | – | 1.74 |
| Weight | – | 69.63 |
| BMI | – | 23 |
| Volume | – | 0.221 |
| Concentration | – | 12 |
| Drink kcal | – | 38.91 |
| Drink length | 10 | – |
| Food kcal | 555 | – |
| Time of meal | -60 | – |

Supplementary Table 24: **Input values for Frezza men**

###### 3.2.25 Kechagias PEth

| Input | Given values | Inferred values |
|---|---|---|
| Sex | – | 0 |
| Height | – | 1.667 |
| Weight | – | 82.1 |
| BMI | 25.0 | – |
| Volume | – | 0.2 |
| Concentration | 13.5 | – |
| Drink kcal | – | 140.9 |
| Drink length | – | 15 |
| Food kcal | 0 | – |
| Time of meal | 0 | – |

Supplementary Table 25: **Input values for Kechagias PEth**

##### 3.3 Calculations for each study and the given values above

###### 3.3.1 Okabe 500ml Water

$$kcal = kcal\_per\_vol \cdot vol = 0 \cdot 0.5 = 0 \text{ kcal} \tag{12}$$

###### 3.3.2 Okabe 500ml orange juice

$$kcal = kcal\_per\_vol \cdot vol = 440 \cdot 0.5 = 220 \text{ kcal} \tag{13}$$

###### 3.3.3 Okabe 500ml milk water

$$kcal = kcal\_per\_vol \cdot vol = 440 \cdot 0.5 = 220 \text{ kcal} \tag{14}$$

###### 3.3.4 Okabe 500ml orange juice and syrup

$$kcal = kcal\_per\_vol \cdot vol = 660 \cdot 0.5 = 330 \text{ kcal} \tag{15}$$

###### 3.3.5 Okabe 500ml milk

$$kcal = kcal\_per\_vol \cdot vol = 660 \cdot 0.5 = 330 \text{ kcal} \tag{16}$$

###### 3.3.6 Okabe 150ml Water

$$kcal\_per\_vol = kcal/vol = 0/0.15 = 0 \text{ kcal/L} \tag{17}$$

###### 3.3.7 Okabe 150ml glucose

$$kcal\_per\_vol = kcal/vol = 67/0.15 = 446.67 \text{ kcal/L} \tag{18}$$

###### 3.3.8 Okabe 150ml whiskey

$$kcal\_per\_vol = kcal/vol = 0/0.15 = 0 \text{ kcal/L} \tag{19}$$

###### 3.3.9 Okabe 200ml unifrom glucose

$$kcal\_per\_vol = kcal/vol = 200/0.2 = 1000 \text{ kcal/L} \tag{20}$$

###### 3.3.10 Okabe 400ml unifrom glucose

$$kcal\_per\_vol = kcal/vol = 200/0.4 = 500 \text{ kcal/L} \tag{21}$$

###### 3.3.11 Okabe 600ml unifrom glucose

$$kcal\_per\_vol = kcal/vol = 200/0.6 = 333.33 \text{ kcal/L} \tag{22}$$

###### 3.3.12 Mitchell

The drinks used in Mitchell et al. contained 0.5 g ethanol per kg body weight [2]. Furthermore, ethanol contains 7 kcal/g and the density of ethanol is 0.7891 g/mL. Using this information we can estimate the total volume of the ethanol in the drink.

$$\begin{aligned}
&height = \sqrt{(}weight/BMI) = \sqrt{(}82.66/226.35) = 1.771 \text{ m}\\
&mass_EtOH = EtOH_dose \cdot weight = 0.5 \cdot 82.66 = 41.33 \text{ g}\\
&EtOH_kcal = mass_EtOH \cdot EtOH_energy = 41.33 \cdot 7 = 289.31 \text{ kcal}\\
&vol_EtOH = EtOH_kcal/density = (41.33/0.7891) \cdot 0.001 = 0.052376 \text{ L}
\end{aligned} \tag{23}$$

**Beer**
For beer, it was reported that a 80 kg male was drinking a total of 409 kcal. From this, we can calculate the total volume, total kcal and total kcal/volume of the drinks, knowing that the reported mean wight was 82.66 kg:

$$\begin{aligned} volume &= (c2 \cdot v2/c1) = (0.052376 \cdot 1)/0.051 = 1.027 \text{ L} \\ total\_kcal &= (kcal/kg) \cdot weight = (409/80) \cdot 82.66 = 422.60 \text{ kcal} \\ Beverage\_kcal &= total\_kcal - EtOH\_kcal = 422.6 - 289.31 = 133.29 \text{ kcal} \\ kcal\_per\_vol &= 133.29/1.027 = 129.79 \text{ kcal/L} \end{aligned} \tag{24}$$

**Wine**
For wine, it was reported that a 80 kg male was drinking a total of 334 kcal. From this, we can calculate the total volume, total kcal and total kcal/volume of the drinks, knowing that the reported mean wight was 82.66 kg:

$$\begin{aligned} volume &= (c2 \cdot v2/c1) = (0.052376 \cdot 1)/0.125 = 0.419 \text{ L} \\ total\_kcal &= (kcal/kg) \cdot weight = (334/80) \cdot 82.66 = 345.11 \text{ kcal} \\ Beverage\_kcal &= total\_kcal - EtOH\_kcal = 345.11 - 289.31 = 55.8 \text{ kcal} \\ kcal\_per\_vol &= 55.8/0.419 = 133.17 \text{ kcal/L} \end{aligned} \tag{25}$$

**Spirit**
For spirit, it was reported that a 80 kg male was drinking a total of 297 kcal. From this, we can calculate the total volume, total kcal and total kcal/volume of the drinks, knowing that the reported mean wight was 82.66 kg:

$$\begin{aligned} volume &= (c2 \cdot v2)/c1 = 0.052376 \cdot 1/0.051 = 0.2618 \text{ L} \\ total\_kcal &= (kcal/kg) \cdot weight = (297/80) \cdot 82.66 = 306.88 \text{ kcal} \\ Beverage\_kcal &= total\_kcal - EtOH\_kcal = 306.88 - 289.31 = 17.57 \text{ kcal} \\ kcal\_per\_vol &= 17.57/0.2618 = 67.11 \text{ kcal/L} \end{aligned} \tag{26}$$

##### 3.3.13 Sarkola

First group of women, female1, has $n = 10$ individuals and anthropometrics, $sex = 0$, $BMI = 21.5$, estimated height 1.66 m as it was average height of women in Finland for the mean age of the participants and year of the study.

$$weight\_female1 = BMI \cdot height^2 = 21.5 \cdot 1.66^2 = 59.24 \text{ kg} \tag{27}$$

Second group of women, female2, has $n = 12$ individuals and anthropometrics, $sex = 0$, $BMI = 21.9$, estimated height 1.66 m as it was average height of women in Finland for the mean age of the participants and year of the study.

$$weight\_female2 = BMI \cdot height^2 = 21.9 \cdot 1.66^2 = 60.35 \text{ kg} \tag{28}$$

The group of men, male, has $n = 13$ individuals and anthropometrics, $sex = 1$, $BMI = 23$, estimated height 1.80 m as it was average height of men in Finland for the mean age of the participants and year of the study.

$$weight\_male = BMI \cdot height^2 = 23.0 \cdot 1.80^2 = 74.52 \text{ kg} \quad (29)$$

**Calculation of a mean person**
We first calculate the blood volumes of the different participant groups using Nadler's equation [3], and the ratio of males in the participant group:

$$\begin{aligned}
vol_blood_female1 &= (0.3561 \cdot height_female1^3 + 0.03308 \cdot weight_female1 + 0.1833) \\
&= 37.718660055999997 \text{ dL} \\
vol_blood_female2 &= (0.3561 \cdot height_female2^3 + 0.03308 \cdot weight_female2 + 0.1833) \\
&= 38.085848056 \text{ dL} \\
vol_blood_male &= (0.3669 \cdot height_male^3 + 0.03219 \cdot weight_male + 0.6041) \\
&= 51.42659600000001 \text{ dL} \\
vol_blood &= (13 \cdot vol_blood_male + 10 \cdot vol_blood_female1 + 12 \cdot vol_blood_female2)/(10 + 12 + 13) \\
&= (13 \cdot 51.42659600000001 + 12 \cdot 38.085848056 + 10 \cdot 37.718660055999997)/(35) \\
&= 42.93607214948572 \text{ dL} \\
sex &= n_male/n_total = 13/(10 + 12 + 13) = 0.37142857142857144
\end{aligned} \quad (30)$$

Since the ratio of males is less than 0.5, we use a female as the average person for this study. We then calculate the weight and height based on the mean blood volume (again using Nadler's equation) to get an approximate woman representing the average person in the study.

$$\begin{aligned}
height &= height_female1 = 1.66 \text{ m} \\
weight &= (vol_blood - 0.1833 - 0.3561 \cdot height^3)/0.03308 = 75.01 \text{ kg} \\
BMI &= weight/height^2 = 75.01/1.66^2 = 27.22\ kg/m^2
\end{aligned} \quad (31)$$

**Estimated inputs**
The drink used by Sarkola et al. contained 0.5 g ethanol per kg body weight [4]. Furthermore, the density of ethanol is 0.7891 g/mL. Using this information we can estimate the total drink volume and kcal per volume.

$$\begin{aligned}
&mass_EtOH = 0.5 \cdot weight = 0.5 \cdot 75.01 = 37.505 \text{ g} \\
&vol_EtOH = mass_EtOH/density = 37.505 \cdot 0.001/0.7891 = 0.047529 \text{ L} \\
&volume = (v1 \cdot c1)/c2 = (0.047529 \cdot 1)/0.1 = 0.475 \text{ L} \\
&Beverage_kcal = (volume - vol_EtOH) \cdot (kcal/L) = (0.475 - 0.047529) \cdot (35/0.1) = 149.61 \text{ kcal} \\
&kcal_per_vol = 149.61/0.475 = 314.97 \text{ kcal/L}
\end{aligned} \quad (32)$$

##### 3.3.14 Jones EtOH and acetaldehyde

The drink used by Jones et al. contained 0.25 g ethanol per kg body weight [5]. Furthermore, the density of ethanol is 0.7891 g/mL. Height was assumed to be 1.79 m as it was the average height of men in Sweden for the mean age of the participants and year of the study. Using this information we can estimate the total drink volume and kcal per volume.

$$\begin{aligned}
&BMI = weight/(height^2) = 74.5/(1.79^2) = 23.25\ kg/m^2\\
&mass_EtOH = 0.25 \cdot weight = 0.25 \cdot 74.5 = 18.625 \text{ g}\\
&vol_EtOH = EtOH_kcal/density = 18.625 \cdot 0.001/0.7891 = 0.023603 \text{ L}\\
&conc = (c1 \cdot v1)/v2 = (0.023603 \cdot 1)/0.250 = 9.44\ \%\\
&Beverage_kcal = (volume - vol_EtOH) \cdot (kcal/L) = (0.250 - 0.023603) \cdot (40/0.1) = 90.56 \text{ kcal}\\
&kcal_per_vol = 90.56/0.250 = 362.24 \text{ kcal/L}
\end{aligned} \tag{33}$$

##### 3.3.15 Jones fasting and food

The drink used by Jones et al. contained 0.3 g ethanol per kg body weight [6]. Furthermore, the density of ethanol is 0.7891 g/mL. Using this information we can estimate the total drink volume and kcal per volume.

$$\begin{aligned}
&BMI = weight/(height^2) = 75.8/(1.83^2) = 22.63\ kg/m^2\\
&mass_EtOH = 0.3 \cdot weight = 0.3 \cdot 75.8 = 22.74 \text{ g}\\
&vol_EtOH = EtOH_kcal/density = 22.74 \cdot 0.001/0.7891 = 0.028818 \text{ L}\\
&volume = (v1 \cdot c1)/c2 = (0.028818 \cdot 1)/0.2 = 0.1441 \text{ L}\\
&Beverage_kcal = (volume - vol_EtOH) \cdot (kcal/L) = (0.1441 - 0.028818) \cdot (45/0.1) = 51.877 \text{ kcal}\\
&kcal_per_vol = 51.877/0.1441 = 360.0 \text{ kcal/L}
\end{aligned} \tag{34}$$

**Food**
The meal was consumed over 15 minutes and approximately 5 additional minutes before the drink was served, which gives $time\_meal = -20$

##### 3.3.16 Kechagias fasting and breakfast

The drink used by Jones et al. contained 0.3 g ethanol per kg body weight [7]. Furthermore, the density of ethanol is 0.7891 g/mL. Height assumed to be 1.80 m as it was average height of men in Sweden for the mean age of the participants and year of the study. Using this information we can estimate the total drink volume and kcal per volume.

$$\begin{aligned}
&BMI = weight/(height^2) = 81.9/(1.80^2) = 25.28\ kg/m^2\\
&mass_EtOH = 0.3 \cdot weight = 0.3 \cdot 81.9 = 24.57 \text{ g}\\
&vol_EtOH = EtOH_kcal/density = 24.57 \cdot 0.001/0.7891 = 0.031137 \text{ L}\\
&volume = (v1 \cdot c1)/c2 = (0.031137 \cdot 1)/0.2 = 0.156 \text{ L}\\
&Beverage_kcal = (volume - vol_EtOH) \cdot (kcal/L) = (0.156 - 0.031137) \cdot (45/0.1) = 56.188 \text{ kcal}\\
&kcal_per_vol = 56.188/0.156 = 360.0 \text{ kcal/L}
\end{aligned} \tag{35}$$

##### 3.3.17 Javors low dose

Height for males assumed to be 1.78 m as it was average height of males in USA for the mean age of the participants and year of the study. Height for females assumed to be 1.63 m as it was average height of females in USA for the mean age of the participants and year of the study.

**Calculation of a mean person**
We first calculate the blood volumes of the different participant groups using Nadler's equation [3], and the ratio of males in the participant group.

$$
\begin{aligned}
&n\_females = 8 \\
&n\_males = 8 \\
&sex = n\_males/(n\_females + n\_males) = 8/(8+8) = 0.5 \\
&vol\_blood\_female = (0.3561 \cdot height\_female^3 + 0.03308 \cdot weight\_female + 0.1833) \\
&\qquad = (0.3561 \cdot 1.63^3 + 0.03308 \cdot 72.7 + 0.1833) \\
&\qquad = 41.3 \text{ dL} \\
&vol\_blood\_male = (0.3669 \cdot height\_male^3 + 0.03219 \cdot weight\_male + 0.6041) \\
&\qquad = (0.3669 \cdot 1.78^3 + 0.03219 \cdot 72.7 + 0.6041) \\
&\qquad = 50.1 \text{ dL} \\
&vol\_blood = (sex) \cdot vol\_blood\_male + (1 - sex) \cdot vol\_blood\_female \\
&\qquad = 0.5 \cdot 50.1 + (1 - 0.5) \cdot 41.3 \\
&\qquad = 45.7 \text{ dL}
\end{aligned}
\tag{36}
$$

Since the ratio of males is equal to 0.5, we used a male as the average person for this study. We then calculate the weight and height based on the mean blood volume (again using Nadler's equation) to get an approximate male representing the average person in the study.

$$
\begin{aligned}
&height = height\_male = 1.78 \text{ m} \\
&weight = (vol\_blood - 0.6041 - 0.3669 \cdot height\_male^3)/0.03219 \\
&\qquad = (4.57 - 0.6041 - 0.3669 \cdot 1.78^3)/0.03219 \\
&\qquad = 58.92124856166512 \text{ kg} \\
&BMI = 58.92124856166512/1.78^2 = 18.60\ kg/m^2
\end{aligned}
\tag{37}
$$

**Estimated inputs**
Knowing that the density of ethanol is equal to 0.7891 g/ml or kg/L and the reported consumption of 0.25 g ethanol per kg bodyweight [1], we can estimate the inputs.

$$
\begin{aligned}
&mass\_EtOH = 0.25 \cdot weight = 0.25 \cdot 72.7 = 18.175 \text{ g} \\
&conc = mass\_EtOH/(density \cdot volume) = (18.175 \cdot 0.001)/(0.7891 \cdot 0.70976) = 3.245\ \% \\
&Beverage\_kcal = (volume - vol\_EtOH) \cdot (kcal/L) = (0.70976 - 0.018175) \cdot (35/0.1) = 242.05 \text{ kcal} \\
&kcal\_per\_vol = 242.05/0.70976 = 341.03 \text{ kcal/L}
\end{aligned}
\tag{38}
$$

###### 3.3.18 Javors high dose

Height for males assumed to be 1.78 m as it was average height of males in USA for the mean age of the participants and year of the study. Height for females assumed to be 1.63 m as it was average height of females in USA for the mean age of the participants and year of the study.

**Calculation of a mean person**
We first calculate the blood volumes of the different participant groups using Nadler's equation [3], and the ratio of males in the participant group.

$$\begin{aligned}
n_females &= 6\\
n_males &= 5\\
sex &= n_males/(n_females + n_males) = 8/(8+8) = 0.545\\
vol_blood_female &= (0.3561 \cdot height_female^3 + 0.03308 \cdot weight_female + 0.1833)\\
&= (0.3561 \cdot 1.63^3 + 0.03308 \cdot 71.8 + 0.1833)\\
&= 41.0 \text{ dL}\\
vol_blood_male &= (0.3669 \cdot height_male^3 + 0.03219 \cdot weight_male + 0.6041)\\
&= (0.3669 \cdot 1.78^3 + 0.03219 \cdot 71.8 + 0.6041)\\
&= 49.8 \text{ dL}\\
vol_blood &= (sex) \cdot vol_blood_male + (1 - sex) \cdot vol_blood_female\\
&= 0.545 \cdot 49.8 + (1 - 0.545) \cdot 41.0\\
&= 45.8 \text{ dL}
\end{aligned} \tag{39}$$

Since the ratio of males is greater than 0.5, we used a male as the average person for this study. We then calculate the weight and height based on the mean blood volume (again using Nadler's equation) to get an approximate male representing the average person in the study.

$$\begin{aligned}
height &= height_male = 1.78 \text{ m}\\
weight &= (vol_blood - 0.6041 - 0.3669 \cdot height_male^3)/0.03219\\
&= (4.58 - 0.6041 - 0.3669 \cdot 1.78^3)/0.03219\\
&= 59.231904044734385 \text{ kg}\\
BMI &= 59.231904044734385/1.78^2 = 18.70\ kg/m^2
\end{aligned} \tag{40}$$

**Estimated inputs**
Knowing that the density of ethanol is equal to 0.7891 g/ml or kg/L and the reported consumption of 0.5 g ethanol per kg bodyweight [1], we can estimate the inputs.

$$\begin{aligned}
&mass_EtOH = 0.5 \cdot weight = 0.5 \cdot 71.8 = 35.9 \text{ g}\\
&conc = mass_EtOH/(density \cdot volume) = (35.9 \cdot 0.001)/(0.7891 \cdot 0.70976) = 6.41\ \%\\
&Beverage_kcal = (volume - vol_EtOH) \cdot (kcal/L) = (0.70976 - 0.0359) \cdot (35/0.1) = 235.85 \text{ kcal}\\
&kcal_per_vol = 235.85/0.70976 = 322.29 \text{ kcal/L}
\end{aligned} \tag{41}$$

###### 3.3.19 Javors combined dose

This dataset is a combination of the low- and high dose data. Where, $n\_lowDose = 16$ and $n\_highDose = 11$. Since the data is a combination of two different data sets, we combined the simulations in the same way. In practice, this results in the following expression (Eq. (42)).

$$simulatedValuesCombined = \frac{16 \cdot simulatedValuesLow + 11 \cdot simulatedValuesHigh}{16 + 11} \quad (42)$$

where simulatedValuesCombined can be e.g. plasma ethanol levels.

###### 3.3.20 Frezza woman

Height for females assumed to be 1.61 m as it was average height of females in Italy for the mean age of the participants and year of the study. BMI for females assumed to be 23 $kg/m^2$. The concentration of the wine was assumed to be 12%.

The drink used by Frezza et al. contained 0.3 g ethanol per kg body weight [8]. Furthermore, the density of ethanol is 0.7891 g/mL. Using this information we can estimate the total drink volume and kcal per volume.

$$\begin{aligned}
&weight = BMI \cdot height^2 = 23 \cdot 1.61^2 = 59.62 \text{ kg} \\
&mass_EtOH = 0.3 \cdot weight = 0.3 \cdot 59.62 = 17.886 \text{ g} \\
&vol_EtOH = mass_EtOH/density = 17.886 \cdot 0.001/0.7891 = 0.02267 \text{ L} \\
&volume = (v1 \cdot c1)/c2 = (0.02267 \cdot 1)/0.12 = 0.1889 \text{ L} \\
&Beverage_kcal = (0.1889 - 0.02267) \cdot (20/0.1) = 33.25 \text{ kcal} \\
&kcal_per_vol = 33.25/0.1889 = 176.02 \text{ kcal/L}
\end{aligned} \quad (43)$$

###### 3.3.21 Frezza men

The drink used by Frezza et al. contained 0.3 g ethanol per kg body weight [8]. Furthermore, the density of ethanol is 0.7891 g/mL. Height for men assumed to be 1.74 m as it was average height of men in Italy for the mean age of the participants and year of the study. BMI for men assumed to be 23 $kg/m^2$. The concentration of the wine was assumed to be 12%. Using this information we can estimate the total drink volume and kcal per volume.

$$\begin{aligned}
&weight = BMI \cdot height^2 = 23 \cdot 1.74^2 = 69.63 \text{ kg} \\
&mass_EtOH = 0.3 \cdot weight = 0.3 \cdot 69.63 = 20.89 \text{ g} \\
&vol_EtOH = mass_EtOH/density = 20.89 \cdot 0.001/0.7891 = 0.02647 \text{ L} \\
&volume = (v1 \cdot c1)/c2 = (0.02647 \cdot 1)/0.12 = 0.221 \text{ L} \\
&Beverage_kcal = (0.221 - 0.02647) \cdot (20/0.1) = 38.91 \text{ kcal} \\
&kcal_per_vol = 38.91/0.221 = 176.1 \text{ kcal}
\end{aligned} \quad (44)$$

###### 3.3.22 Kechagias PEth

Height for males assumed to be 1.805 m as it was average height of males in Sweden for the mean age of the participants and year of the study. Height for females assumed to be 1.667 m as it was average height of females in Sweden for the mean age of the participants and year of the study.

**Calculation of a mean person**

We first calculate the blood volumes of the different participant groups using Nadler's equation [3], and the ratio of males in the participant group.

$$\begin{aligned}
&n_females = 14\\
&n_males = 7\\
&sex = n_males/(n_females + n_males) = 7/(7+14) = 0.3333\\
&weight_male = BMI \cdot height^2 = 25 \cdot 1.805^2 = 81.450625 \text{ kg}\\
&weight_female = BMI \cdot height^2 = 25 \cdot 1.667^2 = 69.472225 \text{ kg}\\
&vol_blood_female = (0.3561 \cdot height_female^3 + 0.03308 \cdot weight_female + 0.1833)\\
&\qquad = (0.3561 \cdot 1.667^3 + 0.03308 \cdot 69.472225 + 0.1833)\\
&\qquad = 41.310416786243 \text{ dL}\\
&vol_blood_male = (0.3669 \cdot height_male^3 + 0.03219 \cdot weight_male + 0.6041)\\
&\qquad = (0.3669 \cdot 1.805^3 + 0.03219 \cdot 81.450625 + 0.6041)\\
&\qquad = 53.836373361125 \text{ dL}\\
&vol_blood = (sex) \cdot vol_blood_male + (1 - sex) \cdot vol_blood_female\\
&\qquad = 0.3333 \cdot 53.836373361125 + (1 - 0.3333) \cdot 41.310416786243\\
&\qquad = 45.48531811265117 \text{ dL}
\end{aligned} \tag{45}$$

Since the ratio of males is less than 0.5, we used a female as the average person for this study. We then calculate the weight and height based on the mean blood volume (again using Nadler's equation) to get an approximate female representing the average person in the study.

$$\begin{aligned}
&height = height_female = 1.667 \text{ m}\\
&weight = (vol_blood - 0.1833 - 0.3561 \cdot height_female^3)/0.03308\\
&\qquad = (4.548531811265117 - 0.1833 - 0.3561 \cdot 1.667^3)/0.03308\\
&\qquad = 82.09284569651804 \text{ kg}
\end{aligned} \tag{46}$$

**Estimated inputs**

Males consumed 2 standard glasses of wine (0.3 L), and females consumed 1 standard glass of wine (0.15 L) [9].

$$\begin{aligned}
&volume = (n_females \cdot vol_females + n_males \cdot vol_males)/(n_females + n_males)\\
&\qquad = (14 \cdot 0.15 + 7 \cdot 0.3)/(7+14)\\
&\qquad = 0.2 \text{ L}\\
&vol_EtOH = volume \cdot conc/density = 0.2 \cdot (0.135/0.7891) = 0.034216 \text{ L}\\
&Beverage_kcal = (volume - vol_EtOH) \cdot (85/0.1) = (0.2 - 0.034216) \cdot 850 = 140.9 \text{ kcal}\\
&kcal_per_vol = 140.9/0.2 = 704.5 \text{ kcal/L}
\end{aligned} \tag{47}$$

### 4 Usage of experimental data

All experimental data were digitalized from the original publications, see references in the tables below. The data was then dived into sets of estimation and validation data, the usage and division of data is highlighted in the tables below. For the estimation data, supplementary table 26, and the validation data, supplementary table 27, the following information is given, 1) the origin of the data, 2) which intervention the data corresponds to, 3) what type of data it is, and 4) how many data points (n) the data set consist of.

| **Study** | **Intervention** | **Type of data** | **n** |
|---|---|---|---|
| Okabe et al. 2015 [10] | Water | Gastric emptying | 4 |
| Okabe et al. 2015 [10] | Orange juice | Gastric emptying | 7 |
| Okabe et al. 2015 [10] | Orange juice & syrup | Gastric emptying | 8 |
| Okabe et al. 2015 [10] | Milk & water | Gastric emptying | 7 |
| Okabe et al. 2015 [10] | Milk | Gastric emptying | 8 |
| Okabe et al. 2017 [11] | Glucose solution 200ml | Gastric emptying | 8 |
| Okabe et al. 2017 [11] | Glucose solution 600ml | Gastric emptying | 11 |
| Okabe et al. 2022 [12] | Whiskey | Gastric emptying | 10 |
| Mitchell et al. 2014 [2] | Beer | BAC | 11 |
| Mitchell et al. 2014 [2] | Wine | BAC | 12 |
| Mitchell et al. 2014 [2] | Spirit blend | BAC | 12 |
| Jones et al. 1997 [13] | Spirit blend | BAC | 13 |
| Jones et al. 1997 [13] | Spirit blend & food | BAC | 13 |
| Kechagias et al. 1999 [7] | Spirit blend | BAC | 20 |
| Kechagias et al. 1999 [7] | Spirit blend & food | BAC | 18 |
| Javors et al. 2016 [1] | Spirit blend high dose | BrAC | 7 |
| Javors et al. 2016 [1] | Spirit blend high dose | PEth | 7 |
| Javors et al. 2016 [1] | Spirit blend combined doses | PEth | 13 |
| Sarkola et al. 2016 [4] | Spirit blend | Acetate | 4 |

Supplementary Table 26: **Estimation data**

| **Study** | **Intervention** | **Type of data** | **n** |
|---|---|---|---|
| Okabe et al. 2017 [11] | Uniform glucose 400ml | Gastric emptying | 10 |
| Okabe et al. 2022 [12] | Water | Gastric emptying | 10 |
| Okabe et al. 2022 [12] | Glucose solution | Gastric emptying | 10 |
| Sarkola et al. 2016 [4] | Spirit blend | BAC | 4 |
| Javors et al. 2016 [1] | Spirit blend low dose | BrAC | 7 |
| Javors et al. 2016 [1] | Spirit blend low dose | PEth | 7 |
| Frezza et al. 1990 [8] | Wine Female | BAC | 13 |
| Frezza et al. 1990 [8] | Wine Male | BAC | 12 |
| Kechagias et al. 2015 [9] | Wine | PEth | 1 |

Supplementary Table 27: **Validation data**
